# A small molecule that inhibits the evolution of antibiotic resistance

**DOI:** 10.1101/2022.09.26.509600

**Authors:** Anna E. Johnson, Harrison Bracey, Angel Joel Hernandez Viera, Juan Carvajal-Garcia, Esra N. Simsek, Kwangho Kim, Houra Merrikh

**Author notes:** These authors contributed equally to this work (Anna E. Johnson, Harrison Bracey).

## Abstract

Antimicrobial resistance (AMR) rapidly develops against almost all available therapeutics. New antibiotics target essential processes in bacteria but fail to address the root of the problem: mutagenesis and evolution. We recently proposed that inhibiting the molecular mechanisms underlying bacterial evolution is the ultimate solution to preventing AMR development. Here, we describe the first compound that inhibits the occurrence and progression of AMR by directly targeting a highly conserved bacterial evolvability factor, Mfd. We previously found that this RNA polymerase-associated translocase is required for rapid AMR development across highly divergent pathogens. Through an *in vivo* screen, we identified 43 potential Mfd-inhibiting compounds. Here we present on target validation, biochemical characterization, and *in vivo* efficacy studies of a lead compound, referred to as ARM-1. ARM-1 binds Mfd and modulates its RNA polymerase interaction. Inhibition of Mfd activity by ARM-1 delays the development of mutations and resistance acquisition, both in pure culture and during infection. Importantly, our data show that this compound prevents the evolution of AMR across highly divergent pathogens, including *Pseudomonas aeruginosa, Staphylococcus aureus, Listeria monocytogenes*, and *Salmonella enterica* serovar Typhimurium. The novel compound we present here has the potential to develop into a clinically useful “anti-evolution” drug. This work demonstrates that the molecular mechanisms of evolution are pharmaceutically targetable, and that this strategy could help prevent AMR development.

## Introduction

Antimicrobial resistance (AMR) is among the top five global health crises of the modern world. World Health Organization priority pathogens such as methicillin-resistant *Staphylococcus aureus* (MRSA) and drug-resistant *Mycobacterium tuberculosis* (DR-TB), along with hundreds of other drug resistant bacterial species, account for global mortality rates over 10 million every year^1^. By 2050, if unchecked, AMR is projected to cost 210 trillion USD in annual global GDP, with roughly 16.7 trillion from DR-TB alone.

Our lab and others have previously proposed a fundamental shift in the approach to combatting AMR^2–5^. Inhibiting evolution by directly inactivating the bacterial mechanisms that increase mutation rates could prevent the development of AMR during the treatment of infections. Our previous findings showed that Mfd, an RNA polymerase (RNAP)-associated DNA translocase, increases mutagenesis and accelerates evolution of resistance across highly divergent species^6–9^. Our work specifically implicates Mfd in acquisition of “spontaneous” point mutations in target genes^9^. Many types of mutagenic processes contribute to the development of resistance, including decreased uptake, enzymatic modification, inactivation, and alterations in the antibiotic target^10^. However, target and nontarget-based mutagenesis (i.e. mutations in genes coding for an antibiotic target or other gene related to resistance acquisition) plays a significant role in resistance development to aminoglycosides^11,12^, glycopeptides^13^, macrolides^11,12^, oxazolidinones^14^, beta-lactams^15,16^, quinolones^17^, tetracycline^11,12^, and chloramphenicol^11^ in diverse species, such as *S. aureus, S. pneumoniae, P. aeruginosa*, and *E. coli*. In this work, we describe our discovery of a novel small molecule that modulates the activity Mfd, demonstrating both that directly targeting bacterial evolvability factors is possible and can impede the development of AMR by reducing target and nontarget-based mutagenesis.

## Results

### An *In Vivo* High-Throughput Screen Identified Novel Mfd-Inhibiting Compounds

To determine if Mfd activity can be inactivated through the use of a small molecule, we conducted a high-throughput *in vivo* screen of roughly 250,000 novel compounds (S.I. Fig. 1). We modified the previously described roadblock repression assay to reveal Mfd-specific inhibitors in a biologically relevant context (Fig. 1a, S.I. Fig. 1)^18^. We have previously shown that interactions of Mfd with RNAP and the nucleotide excision repair protein UvrA are required for its pro-mutagenic effects^6^; assaying potential inhibitor compounds *in vivo* maintains these interactions and allows us to discover compounds that modulate Mfd’s physical interactions as well as its enzymatic activity. A further advantage of this design is that it will only identify small molecules that are able to permeate bacterial cells and have little or no bactericidal activity or toxicity to mammalian cells. We used Δ*mfd Escherichia coli* cells containing a reporter system encoded on two different plasmids (see STAR Methods). One plasmid contains a *lac* operator (*lacO*) encoded directly upstream of the *lux* operon, which expresses proteins that produce luciferase, leading to readily detectable luminescence. The second plasmid expresses the *Salmonella enterica* serovar Typhimurium ST19 Mfd protein under the control of an IPTG-inducible promoter. The LacI protein, which binds to *lacO*, is a barrier to transcription; RNAP is unable to readily pass the bound protein and the transcription machinery will stall. The stalled RNAP is quickly recognized by Mfd and is pushed off the DNA. However, LacI “breathes” on and off DNA at a significant rate^19^. If Mfd is not present, or its activity is inhibited by a small molecule, the stalled RNAP will be able to proceed through the *lux* operon during the temporary dissociation of LacI from the operator sequence, leading to the production of luminescence (Fig. 1a). Using this property, we measured relative luminescence in the presence of each compound to look for Mfd inhibition in this *in vivo* screen (Fig. 1a, S.I. Fig. 1). To control for off target effects, we assayed strains carrying plasmids that did not contain the *mfd* gene or coding for Mfd L499R mutants that are unable to interact with RNAP and examined transcription-dependence of the observed effects in parallel with the experimental conditions. Positive hits from this system are those which result in high luminescence only when transcription is induced and Mfd is present, and do not show prohibitive bactericidal or cytotoxic effects (S.I. Fig. 1, 6). This approach yielded 43 hits. Here we present our investigation of the biochemical and biological effects of the lead compound, VU001, hereafter referred to as ARM-1 (Anti-Resistance Molecule 1). The structure of ARM-1 (VU001) is shown in Fig. 1b.

**Figure 1:**
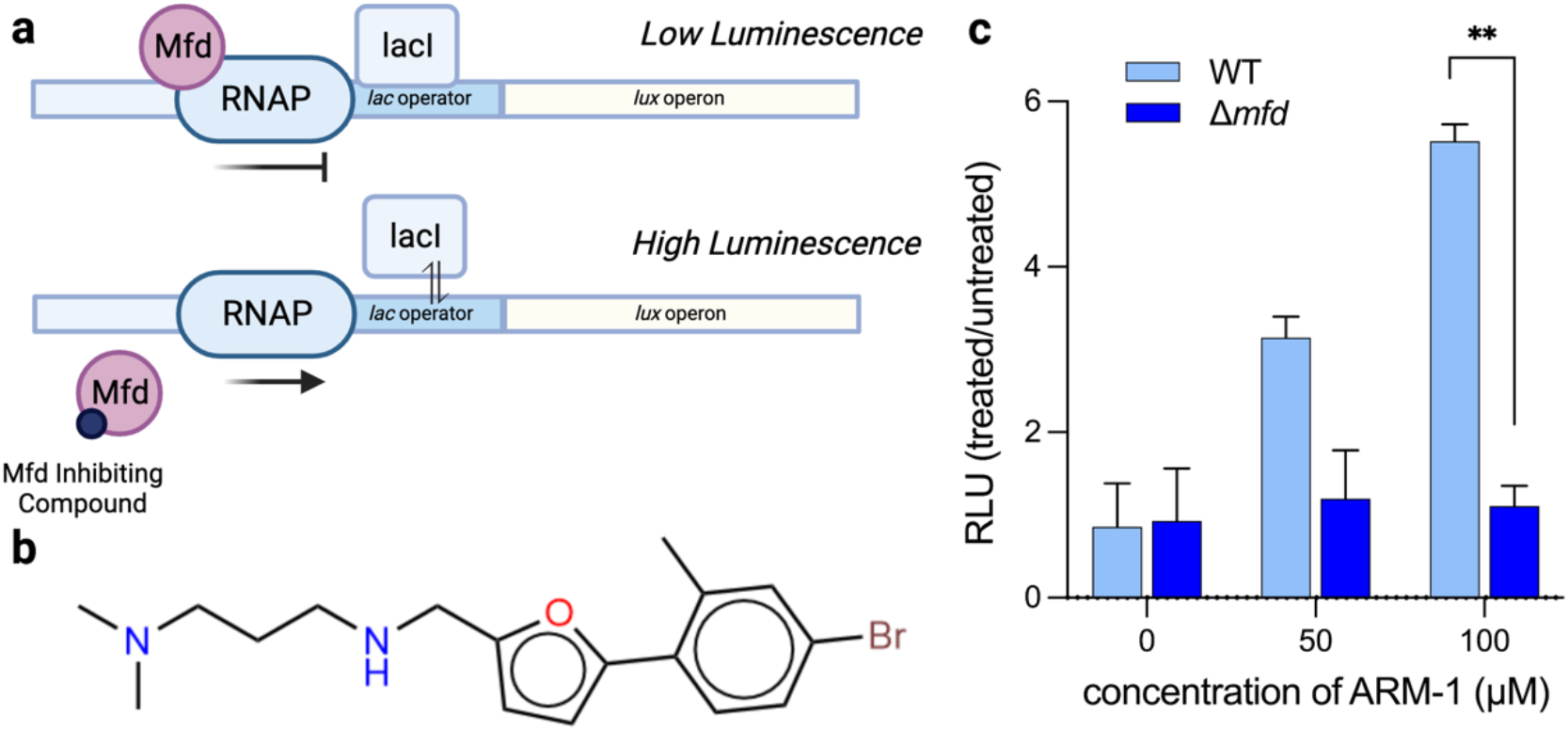
*In vivo* high throughput screen identifies lead compound ARM-1. **a** Schematic of *in vivo* screen design. In the upper panel, when Mfd is present, RNAP complexes stalled at the lacI-bound operator sequence are quickly removed, resulting in little to no transcription of the *lux* operon and low luminescence output. In the lower panel, when Mfd is either absent or inhibited, RNAP complexes stalled at the operator are not removed and are able to proceed during temporary dissociation of LacI, resulting in relatively high levels of luminescent output. **b** Structure of the lead compound, ARM-1. **c** Translocase activity assay showing relative luminescent output for WT and 11*mfd S. enterica* ST19 strains containing the roadblock reporter assay system with increasing concentrations of ARM-1. Relative luminescence is normalized to no compound and 0.5% DMSO solvent control. Statistical significance determined via Welch’s t-test, *p* = 0.005921. ** *p* < 0.01

We synthesized ARM-1 to 299% purity following an original synthesis strategy (see STAR Methods and Supplementary Information for synthesis description and compound validation) and repeated the experiments performed in the screen prior to further investigation into the compound’s effects on Mfd function. We found that ARM-1 binds to Mfd with a k_d_ of 4.25μM (Fig. 2a), an acceptably tight binding affinity for a lead compound prior to medicinal chemistry optimization. This allowed us to use relatively low concentrations of ARM-1 in *in vivo* assays, effectively circumventing any potential bactericidal or cytotoxic effects. As the reaction conditions and quantity of Mfd in each of the experiments presented here are optimized for the individual assays, the amount of ARM-1 was also adjusted to maintain comparable treatment conditions across all experiments (see Table 6). We measured and confirmed the effect of ARM-1 on Mfd’s translocase activity using by introducing the roadblock reporter assay into *S. enterica* ST19. Treating cells with 50 to 100 μM ARM-1 resulted in a 3-to 5-fold inhibition of translocase activity *in vivo*. Given that there was no effect of ARM-1 on luminescence in cells lacking Mfd, we conclude that the effects of ARM-1 in this assay are directly through inhibition of Mfd (Fig. 1c).

**Figure 2.**
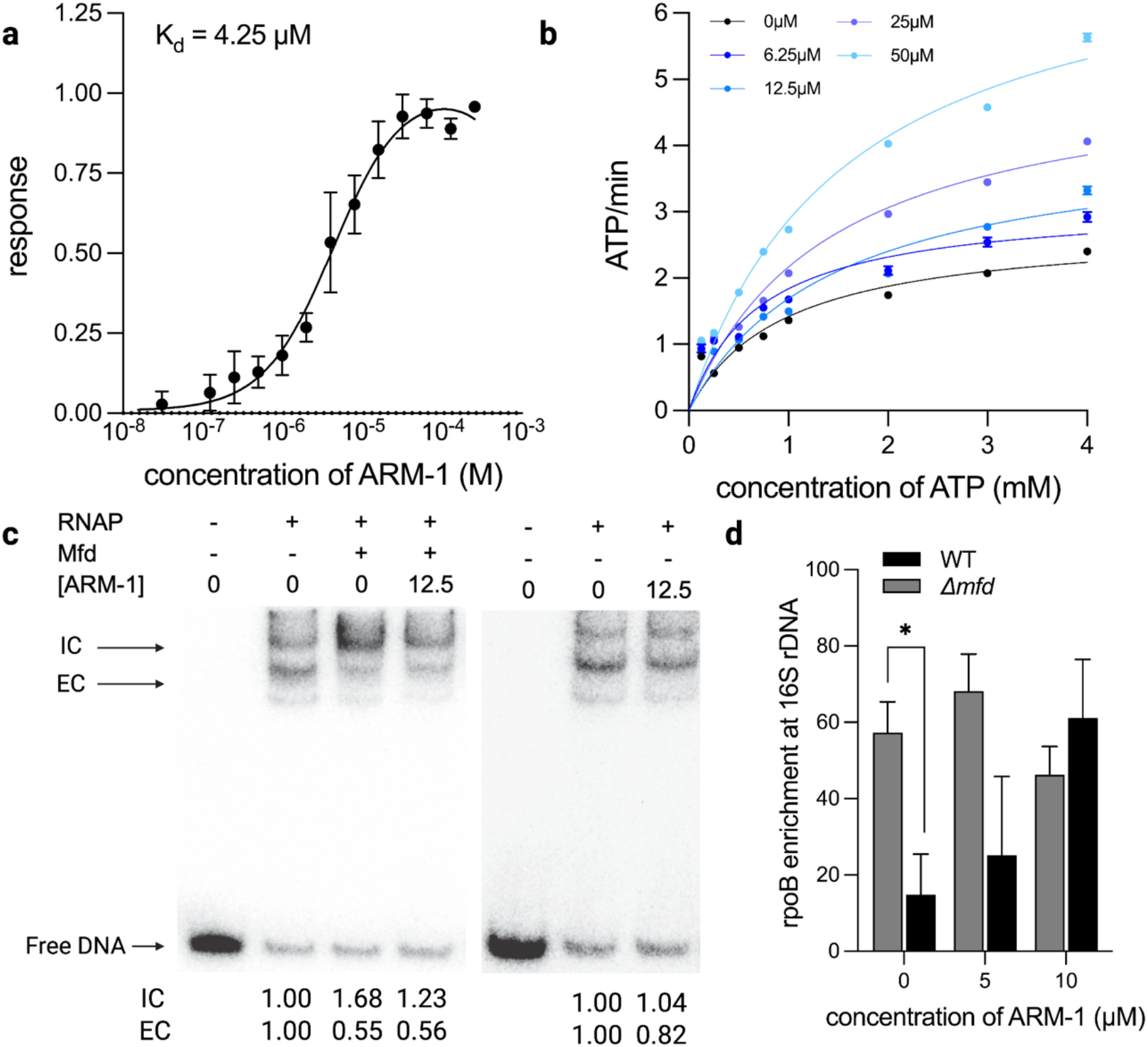
ARM-1 modulates Mfd’s enzymatic activity and interaction with RNAP. **a** k_d_ of ARM-1 was determined using microscale thermophoresis. Data show represent a minimum of three independent experiments. **b** NADH-coupled ATPase assay^35^. ATP hydrolysis by Mfd was coupled to NADH oxidation. Phosphoenolpyruvate is converted to pyruvate by pyruvate kinase, transferring a phosphate group onto ADP, which is only present in the reaction following ATP hydrolysis by Mfd. Pyruvate is then converted to lactate by lactate dehydrogenase, oxidizing NADH to NAD^+^. Absorbance at 340nm by NADH is used to monitor the reaction. Data shown represent a minimum of 3 biological replicates. **c** Transcription roadblock assay. A ^32^P-labled 176bp PCR fragment was incubated with *E. coli* RNAP along with saturating concentrations of ATP, GTP, and UTP. 100nM *S. enterica* ST19 Mfd preincubated with indicated concentrations of ARM-1 was added and the reaction allowed to proceed for 6 minutes. DNA products were then resolved on a polyacrylamide gel and analyzed by phosphorimaging. On the left gel, Lane 1 shows no enzyme and no compound control. Lanes 2 and 3 show RNAP alone and RNAP with Mfd, respectively, both in the absence of ARM-1 treatment. Lanes 4 shows RNAP and Mfd in the presence 12.5μM ARM-1. On the right gel, Lane 1 shows the no enzyme and the no compound control, Lane 2 shows RNAP alone, and Lane 3 shows RNAP with 12.5μM ARM-1. Quantification of these gels is shown below each lane. Analysis was performed using Image Lab 6.0.1. Data shown representative of at least two separate experiments (see Supplemental Information Figure 3). **d** ChIP-qPCR of RpoB at 16S rDNA. *S. enterica* ST19 of indicated genotype grown to mid exponential phase in cells treated with indicated concentration of ARM-1 for 2 generations prior to harvest. ChIP was performed using 8RB13 monoclonal antibody against RpoB. Data shown are representative of 3 biological replicates. Statistical significance determined via Welch’s t-test, p = 0.0459. * *p* < 0.05

### ARM-1 Modulates Mfd’s Enzymatic Activity and Interaction With RNA Polymerase

Mfd translocates along double-stranded DNA and dislodges stalled RNAP complexes^20,21^. As both processes are dependent on Mfd’s ability to hydrolyze ATP, we directly examined the effect of ARM-1 on *in vitro* ATP hydrolysis by Mfd^22,23^. At dosages ranging from 6.25μM to 50μM ARM-1, at molar ratios of 125 to 1000 ARM-1 to Mfd, we observe a dose dependent increase ATP hydrolysis (Fig. 2b, Table 1). ATPase activity is enhanced almost 3-fold in the presence of 50μM ARM-1 (Fig. 2b, Table 1). However, it is important to note that the relative amount of ARM-1 required for any significant effect on ATP hydrolysis by Mfd is far higher than is biologically relevant. At concentrations of ARM-1 consistent with our *in vivo* assays, discussed below, we do not observe any effect on ATP hydrolysis (see Table 6 for comparison of ARM-1 concentrations and ARM-1 to Mfd molar ratios used in each assay). Therefore, we do not expect that a direct effect on ATP hydrolysis is a primary avenue via which ARM-1 modulates Mfd activity.

**Table 1.** Kinetic constants of Mfd ATPase activity determined using Michaelis-Menten modeling. Related to Figure 2b.

| ARM-1 Treatment | Vmax |  | km |  |
| --- | --- | --- | --- | --- |
|  | mean | SEM | mean | SEM |
| 0 μM | 2.771 | 0.343 | 0.954 | 0.319 |
| 6.25 μM | 3.135 | 0.338 | 0.706 | 0.228 |
| 12.5 μM | 4.170 | 0.711 | 1.470 | 0.583 |
| 25 μM | 5.216 | 0.631 | 1.408 | 0.403 |
| 50 μM | 7.401 | 0.666 | 1.576 | 0.322 |

To further investigate the mechanism by which ARM-1 affects Mfd activity, we performed a slightly modified version of an established *in vitro* RNA polymerase displacement assay^24^. In this assay, *E. coli* RNAP holoenzyme is stalled by CTP starvation on a short, radiolabeled DNA fragment containing a constitutively active promoter (see STAR Methods). The first guanine encoded on this DNA fragment is 21 nucleotides downstream of the promoter. In the absence of CTP, RNAP stalls at this location. This stalled RNAP can then be removed by Mfd. Performing an electrophoretic mobility shift assay (EMSA) with the products of this transcription roadblock reaction reveals two DNA-protein complexes: a slower migrating initiation complex (IC), and a faster migrating elongation complex (EC)^24,25^. When quantified, we observed a ratio of EC to IC of consistent with prior work^23,26,27^. Addition of purified Mfd removes the stalled RNA polymerase, which results in a reduction of the ratio of ECs compared to ICs. We reason that when removed by Mfd, the displaced RNAP is free to re-bind the promoter, which is apparent by the increase in the amount of ICs (Fig. 2c, S.I. Fig. 3-4). In contrast, when we pre-incubated Mfd with ARM-1, though we still observed a decrease in the amount of EC when compared to the reaction without Mfd, we no longer observed the corresponding increase in ICs (Fig. 2c, S.I. Fig. 3-4), which is normally a downstream result of Mfd’s RNAP removal function. This indicates that ARM-1 inhibits Mfd’s function by interfering with its ability to remove stalled RNAPs from DNA, notably at a concentration that does not affect its ATP hydrolysis activity. The reduction in the ECs in the presence of ARM-1 is most likely because, at equilibrium, when RNAP cannot dissociate from DNA, it cannot re-initiate transcription, consequently preventing new EC formation. Importantly, the effects of ARM-1 in these experiments are not due to off target effects on RNAP. In our control reactions, ARM-1 does not have a significant effect on the *in vitro* activity RNAP in the absence of Mfd (Fig. 2c). This suggests that our observations are most likely due to Mfd-dependent activity of ARM-1, rather than off-target effects on RNAP. Considering these results together with the ATPase activity assays, we propose that Mfd may displace RNAP not only through enzymatic activity, but also through an allosteric mechanism, which is common to transcription terminators such as Rho^28,29,30^.

We next investigated the effect of ARM-1 on Mfd *in vivo*. For this, we performed chromatin immunoprecipitations (ChIPs) of the β subunit of RNAP (RpoB) in WT and Δ*mfd S. enterica* ST19 strains, treated with ARM-1 for at least two generations of exponential growth. In the untreated condition, we observe an increase in RpoB association with rDNA, a highly transcribed region of the bacterial genome, in the absence of Mfd (Fig. 2d, S.I. Fig. 5). This effect is consistent with Mfd’s function in displacing stalled elongation complexes, particularly at regions that are difficult to transcribe^31^. At 5μM ARM-1, there is little change in RpoB association, but at 10μM there is a dramatic increase in association of RpoB in WT cells, consistent with Mfd inhibition and extended occupancy of elongation complexes at the target region (Fig. 2d, S.I. Fig. 5). In the Δ*mfd* background, contrary to our expectations, RpoB enrichment is not significantly different from WT in the presence of ARM-1. This suggests that *in vivo* ARM-1 may have a limited effect on elongating RNAP complexes independent of Mfd (Fig. 2c, *right panel*, Fig. 2d). Because we observe no growth defects at these concentrations of ARM-1 (S.I. Fig. 7), these patterns are unlikely to be due to indirect effects negatively impacting RNAP activity. In summary, these data support a nuanced effect of ARM-1 on Mfd’s biochemical activity both *in vitro* and *in vivo*.

### ARM-1 Treatment Reduces Mutation Frequency in Culture and During Infection

Our lab has previously shown that the mutagenic effect of Mfd is conserved during bacterial infection of eukaryotic cells. Luria-Delbruck assays have demonstrated that mutation rates decrease in the absence of Mfd^6^. With ARM-1 treatment, we observe a 3-fold reduction in WT *S. enterica* ST19 mutation frequency, consistent with direct inhibition of Mfd (Fig. 3a). To determine if ARM-1 can also reduce mutagenesis during infection, we infected HeLa cells with wild-type and Δ*mfd S. enterica* ST19, and measured mutation frequency through acquisition of 5-fluorocytosine resistance^6,32^. 31.25μM ARM-1 was added to the culture media following to ensure uniform treatment conditions. This treatment condition was determined empirically to avoid any toxicity to HeLa cells and still have a significant impact on bacterial mutation frequency (S.I. Fig. 6). We observed an almost 7-fold decrease in mutation frequency in the absence of Mfd (Fig. 3c). WT *S. enterica* cells have a mutation frequency of 22.3 mutations per 10^5^ bacteria during infection, whereas Δ*mfd* strains have a 3.3 mutations per 10^5^ bacteria during infection (Fig. 3c). We found that treatment with ARM-1 reduces WT mutation frequency to 3.2 mutations per 10^5^ bacteria, a 7-fold reduction compared to untreated cells, which were consistent with rates we observe for Δ*mfd* strains (Fig. 3c). Neither the Δ*mfd* mutant nor ARM-1 or solvent-treated WT or Δ*mfd* strains show a defect in invasion or proliferation of HeLa cells (Fig. 3b). These observations confirm that the differences in mutation frequencies during infection are not due to altered growth dynamics. Taken together, these data strongly suggest that ARM-1 is directly inhibiting bacterial mutagenesis mechanisms during infection.

**Figure 3.**
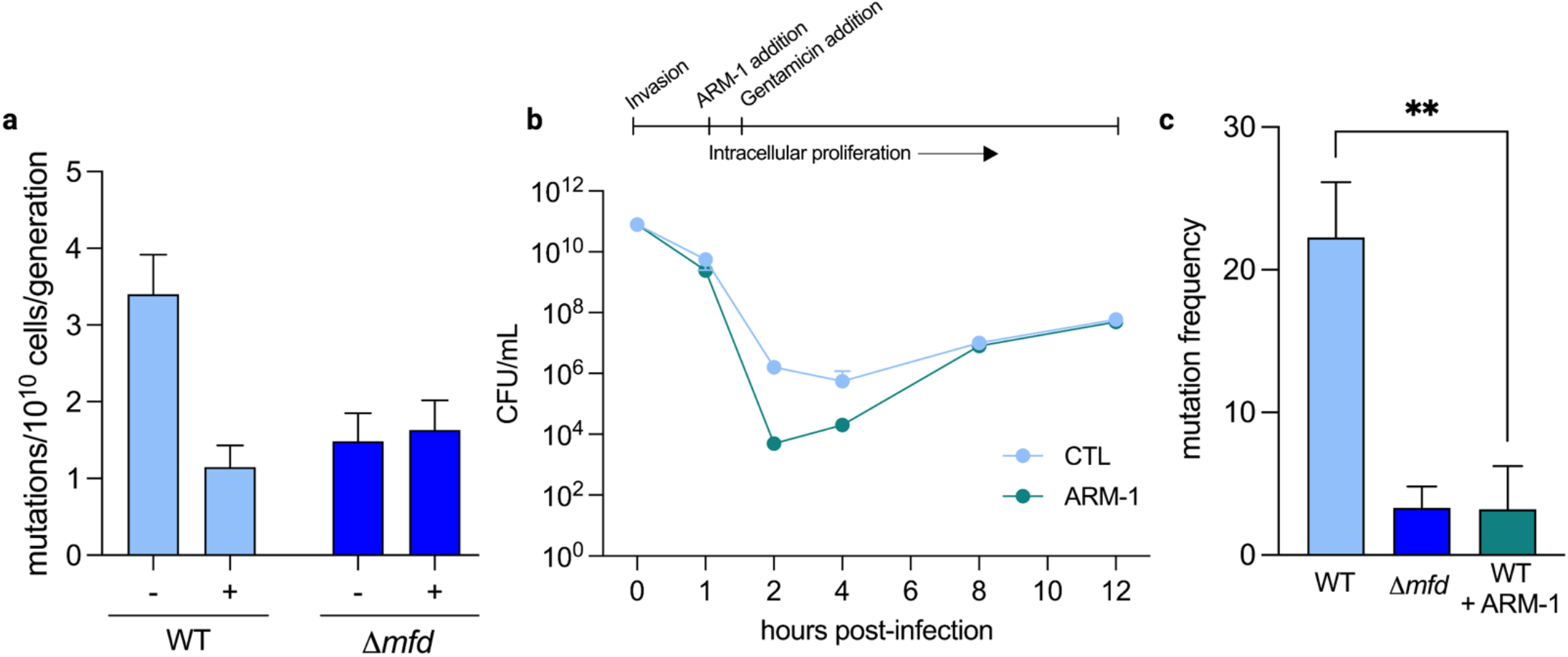
ARM-1 reduces mutation frequency in culture and during infection. **a** Luria-Delbruck fluctuation assay with ARM-1 treatment. Single colonies of WT and Δ*mfd S. enterica* ST19 were used to inoculate overnight cultures with appropriate antibiotic selection. Cultures were then diluted back to OD_600_ 0.0005 and grown to OD_600_ 0.5 with or without 100μM ARM-1 treatment. Samples were plated on 80ng/ml ciprofloxacin and grown overnight, then enumerated the following day. Data shown are the result of a minimum of 50 replicates. **b** Invasion and infection efficiency in the presence of ARM-1. WT *S. enterica* ST19 in mid-log phase growth was used to infect HeLa cells. Invasion was allowed to proceed for 1 hour before uninternalized bacteria were removed and fresh media applied, containing vehicle control or 31.25μM ARM-1. At the indicated timepoints, HeLa cells were lysed using 1% Triton-H_2_O and intracellular bacteria serially diluted and plated on LB for CFU enumeration. **c** Mutation frequency post mammalian cell infection. WT and Δ*mfd S. enterica* ST19 were used to infect HeLa cells as described in **b** Following infection, intracellular bacteria were harvested and plated on 50μg/mL 5-fluorocytosine and grown overnight. Mutants were enumerated the following morning. Mutation frequency is shown as mutants per 10^5^ bacteria harvested from HeLa lysates. Data shown are the result of a minimum of 3 biological replicates. Statistical significance determined via Welch’s t-test, p = 0.0082. ** *p* < 0.01

### ARM-1 Inhibits Development of Antibiotic Resistance

We next examined the effect of ARM-1 on the evolution of antibiotic resistance. Genetic deletion of Mfd is known to reduce the rates and extent of resistance development in laboratory evolution experiments^6,33^. We assessed the impact of ARM-1 on resistance development across diverse pathogens and in response to multiple classes of antibiotics. In these experiments, bacteria are challenged with increasing concentrations of antibiotic over a minimum of 55 generations. We observed a significant reduction in resistance development when bacteria were treated with ARM-1 compared to those that were untreated, in all species and antibiotics tested. This effect was consistent even in strains of bacteria already carrying resistance mutations to other antibiotics, specifically in patient isolates of *S. enterica* ST19 (U.S. patient isolate collected 2000-2010^34^), *S. aureus*, and *P. aeruginosa*. In *S. enterica* ST19, challenged with rifampicin or trimethoprim, we observed at minimum a 100-fold reduction in median minimum inhibitory concentration (MIC) between untreated and treated conditions in these experiments (Fig. 4a-b). This pattern was also observed in *P. aeruginosa, S. aureus*, and *L. monocytogenes*, with ARM-1 treatment resulting in 80, 800, and 1000 reduction in resistance to rifampicin, respectively (Fig. 4a). We observed very similar results when these bacteria were treated with trimethoprim (Fig. 4b). Doubling time was consistent between treated and untreated population across all species tested (S.I. Fig. 7), indicating that our results are not simply attributable to growth defects. Sequencing of relevant resistance loci over the course of treatment confirmed that the reduced rate of resistance development is related to point mutations in target and nontarget genes (Tables 2-5). Populations of cells that were treated with ARM-1 accumulated significantly less mutations compared to untreated cells over the course of the experiment (Tables 2-5). As expected, given the many routes via which bacteria acquire resistance, not all increases in MIC were accompanied by a related target mutation; however, considering the significant reduction in resistance development in ARM-1 treated populations, this result further confirms mutagenesis as a significant factor in resistance acquisition.

**Table 2.**
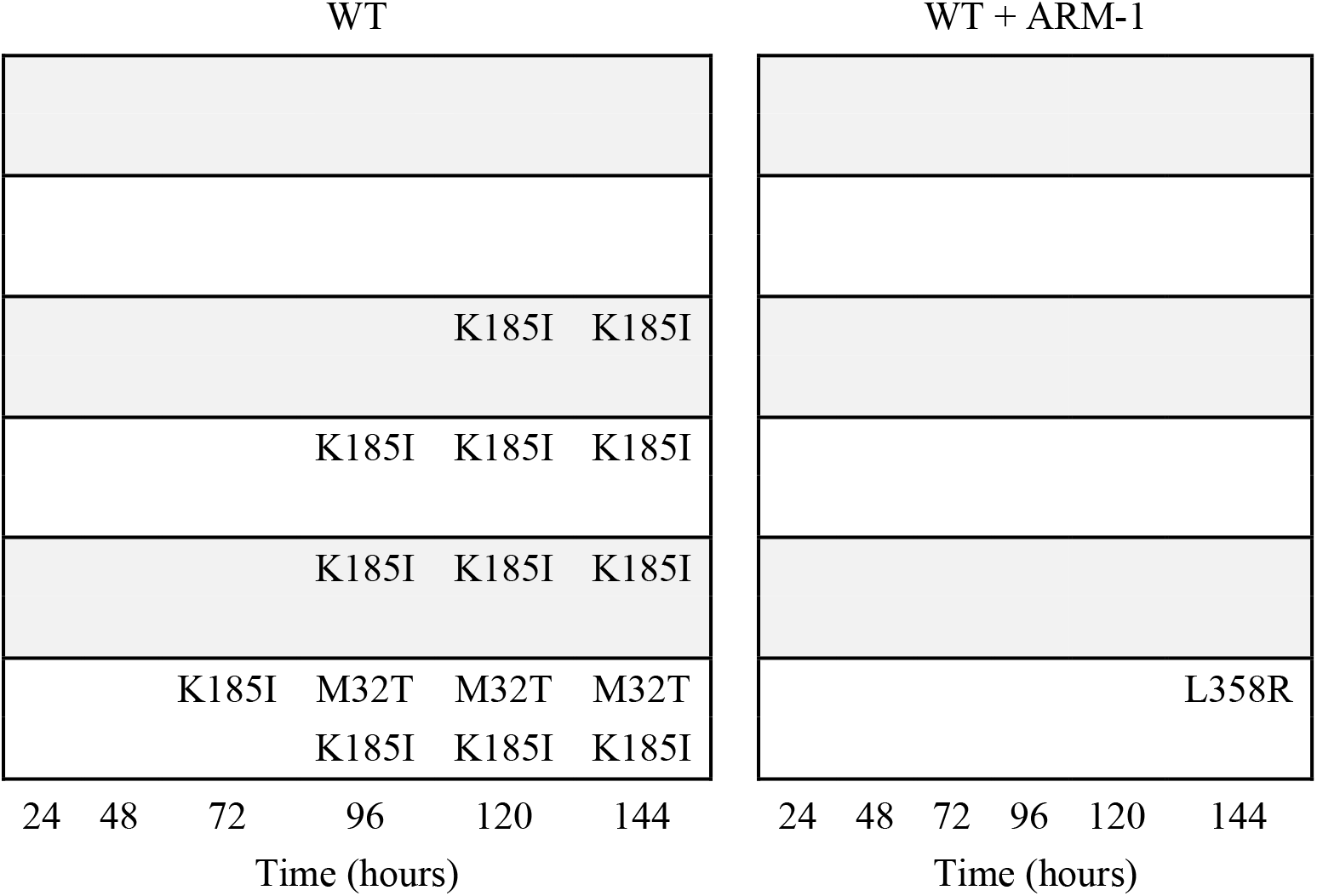
Resistance mutations arising in S. aureus against trimethoprim, with and without ARM-1 treatment. *folA* resistance loci sequenced in 6 biological replicates over the course of an evolution assay. The sample with the highest MIC was selected from each biological replicate. Blank spaces indicate that no mutations were detected in the locus of interest at that timepoint.

**Table 3.**
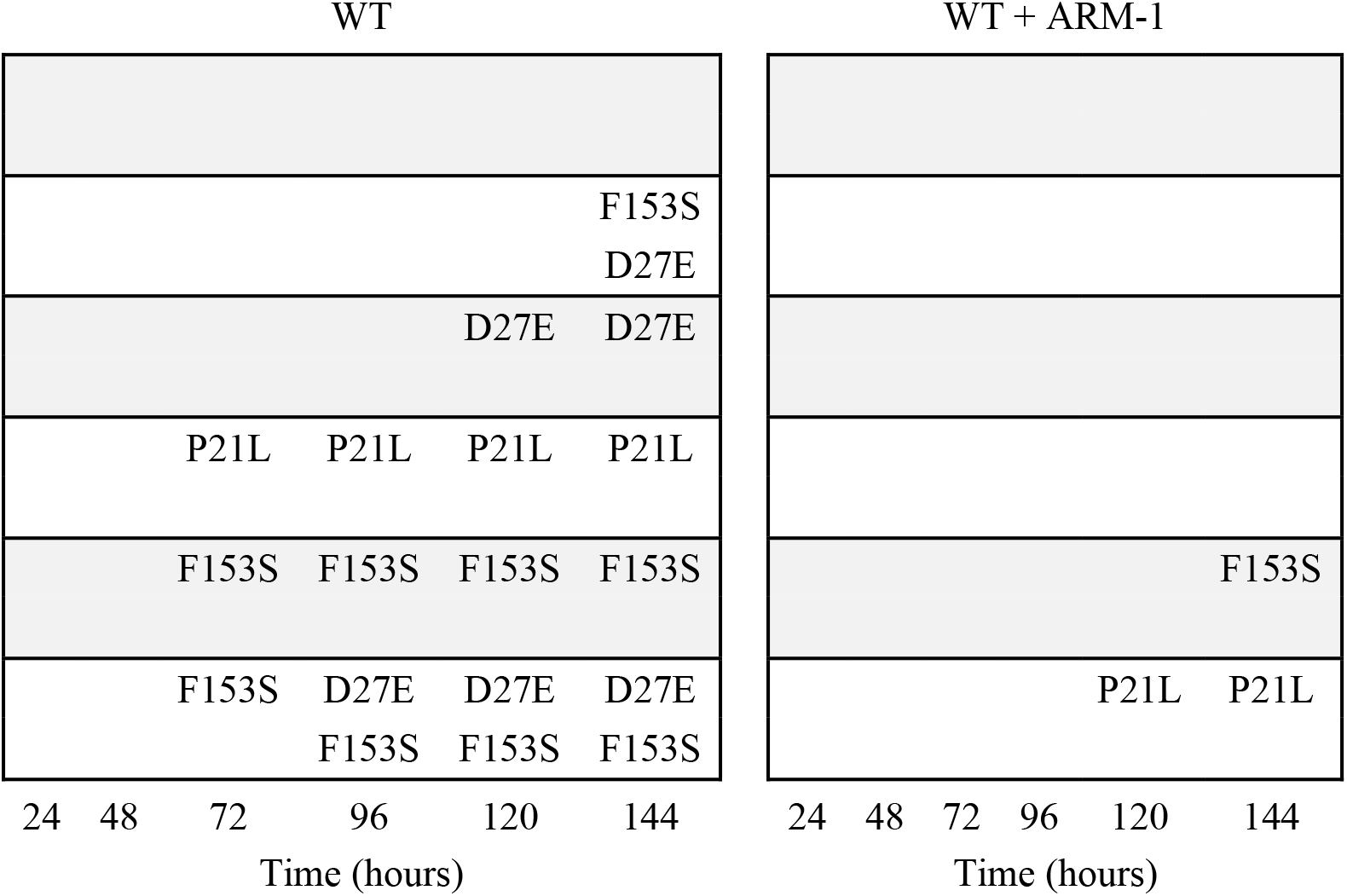
Resistance mutations arising in S. enterica serovar Typhimurium ST19 against trimethoprim, with and without ARM-1 treatment. *folA* resistance loci sequenced in 6 biological replicates over the course of an evolution assay. The sample with the highest MIC was selected from each biological replicate. Blank spaces indicate that no mutations were detected in the locus of interest at that timepoint.

**Table 4.**
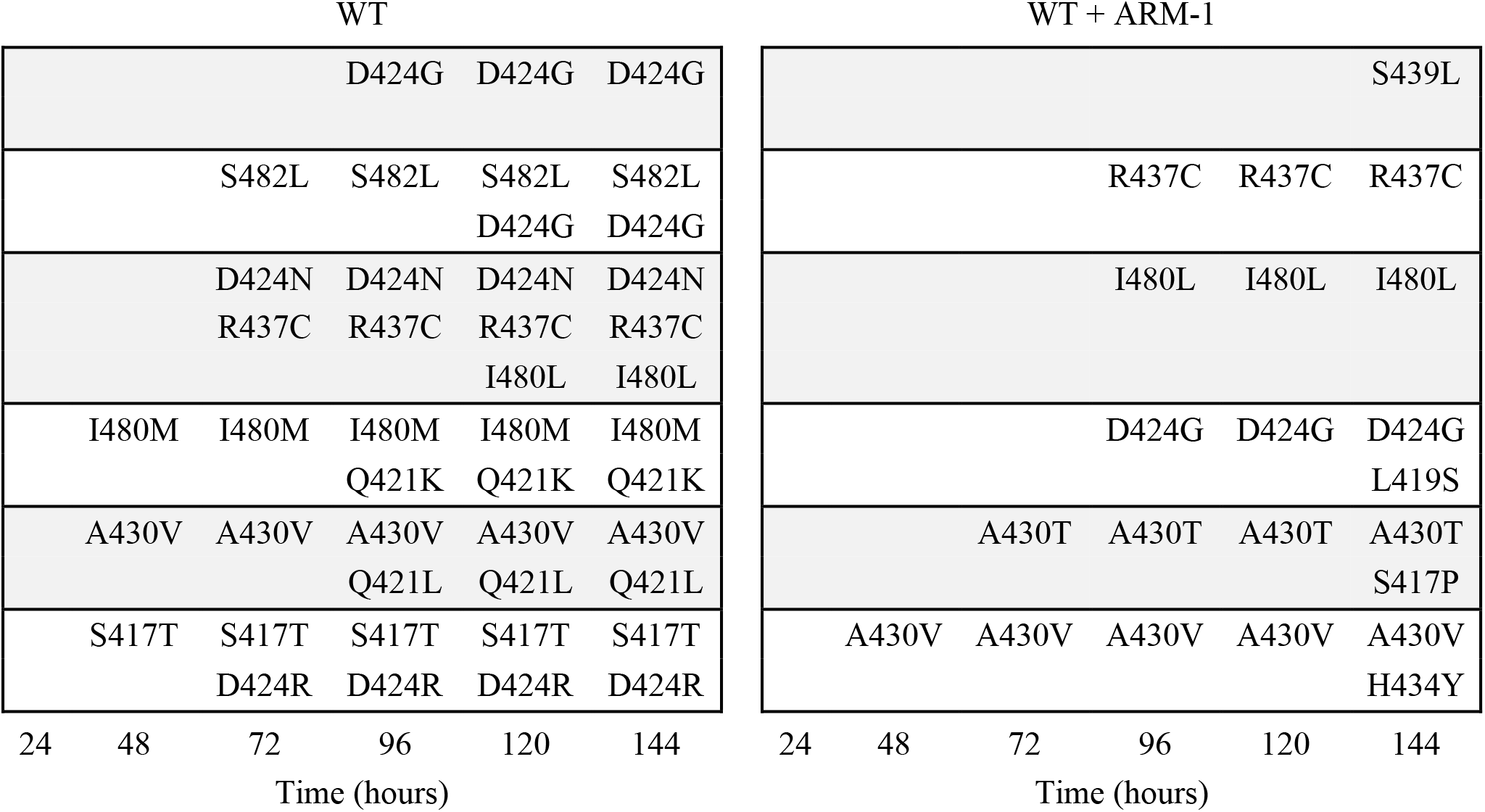
Resistance mutations arising in S. aureus against rifampicin, with and without ARM-1 treatment. *rpoB* RRDR resistance loci sequenced in 6 biological replicates over the course of an evolution assay. The sample with the highest MIC was selected from each biological replicate. Blank spaces indicate that no mutations were detected in the locus of interest at that timepoint.

**Table 5.**
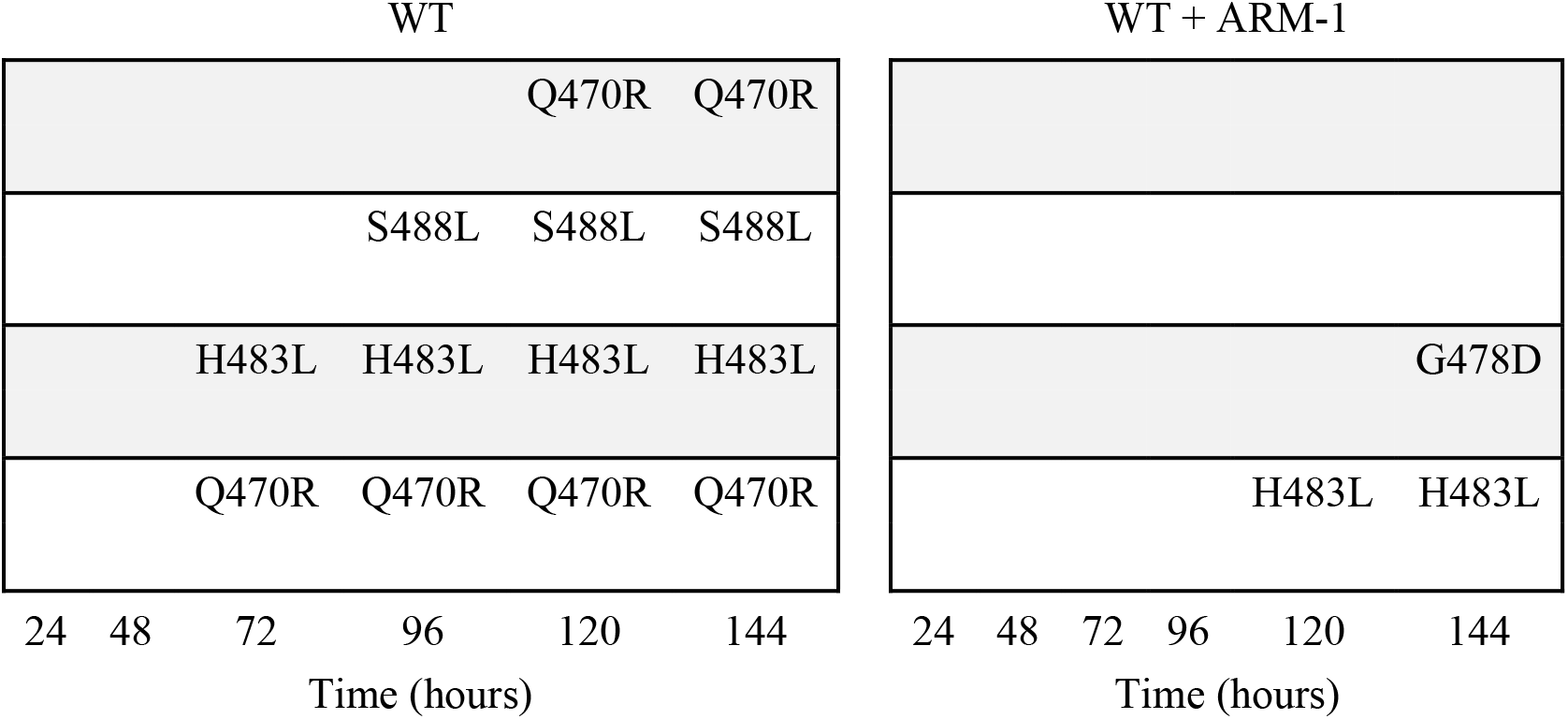
Resistance mutations arising in L. monocytogenes against rifampicin, with and without ARM-1 treatment. *rpoB* RRDR resistance loci sequenced in 6 biological replicates over the course of an evolution assay. The sample with the highest MIC was selected from each biological replicate. Blank spaces indicate that no mutations were detected in the locus of interest at that timepoint.

**Table 6.** Comparison of Relative ARM-1 to Target Molarities by Assay. \**In vivo* assays include evolution assays, rpoB chromatin immunoprecipitations, and infections **Minimum [ARM-1] is lowest concentration at which an effect on RNAP binding is observed in ChIP experiments ***Maximum [ARM-1] is the highest concentration used in in vivo assays, including evolution assays

| MST Binding Affinity |  | Ratio Mfd : ARM-1 |
| --- | --- | --- |
| [Mfd] (M) | 5.00E-09 |  |
| ARM-1 kd | 4.25E-06 | 1 Mfd : 850 ARM-1 |
| [ARM-1] (M) for maximum binding | 1.00E-04 | 1 Mfd : 20,000 ARM-1 |

| ATPase Activity |  | Ratio Mfd : ARM-1 |
| --- | --- | --- |
| [Mfd] (M) | 5.00E-08 |  |
| [ATP] (M) (max) | 4.00E-03 |  |
| Maximum [ARM-1] (M) | 5.00E-05 | 1 Mfd : 1000 ARM-1 |
| Minimum [ARM-1] (M) | 6.25E-06 | 1 Mfd : 125 ARM-1 |

| RNAP Displacement |  | Ratio of Mfd : ARM-1 | Ratio RNAP : ARM-1 |
| --- | --- | --- | --- |
| [Mfd] (M) | 1.00E-07 |  |  |
| [RNAP] (M) | 5.00E-09 |  |  |
| Minimum [ARM-1] (M) for effect | 1.25E-05 | 1 Mfd : 125 ARM-1 | 1 RNAP : 2500 ARM-1 |

| *In vivo* Assays* |  | Ratio of Mfd : ARM-1 | Ratio RNAP : ARM-1 |
| --- | --- | --- | --- |
| [Mfd] (M) per cell | 2.54E-04 |  |  |
| [RNAP] (M) per cell | 2.54E-03 |  |  |
| Minimum [ARM-1] (M)** | 2.50E-05 | 25 Mfd : 1 ARM-1 | 250 RNAP : 1 ARM-1 |
| Maximum [ARM-1] (M)*** | 1.00E-04 | 2.5 Mfd : 1 ARM-1 | 25 RNAP : 1 ARM-1 |

**Figure 4.**
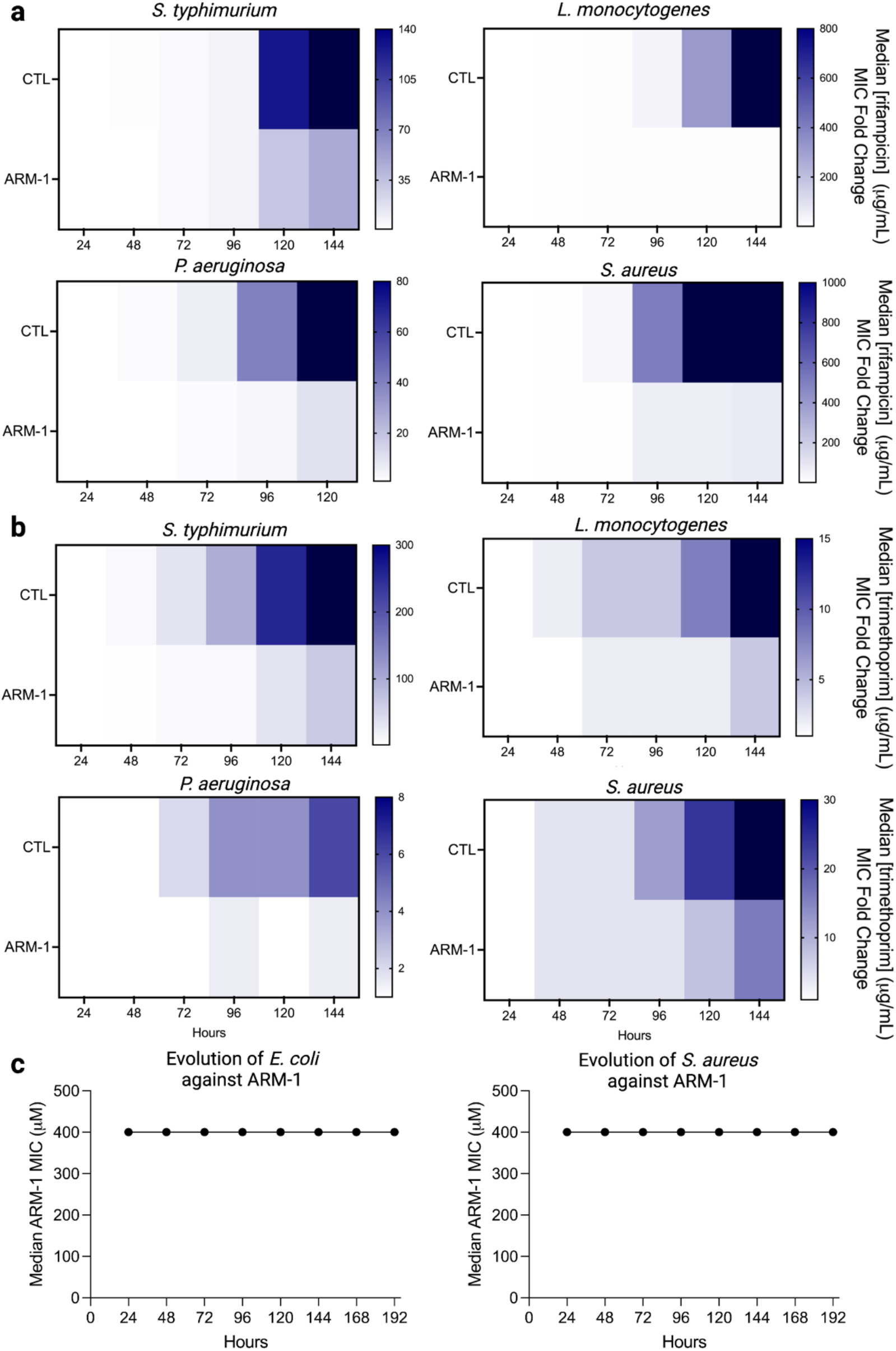
ARM-1 inhibits the evolution of antibiotic resistance. **a-b** Evolution of indicated species against **a** rifampicin and **b** trimethoprim in the presence of ARM-1. Heatmaps show median MIC over time. Concentrations are indicated to the right of each plot. Data for strain/antibiotic combination presented is the result of at least 12 biological replicates. Concentration of ARM-1 used against *L. monocytogenes, P. aeruginosa*, and *S. aureus* is 100μM. Concentration of ARM-1 used against *S. enterica* is 50μM. Populations of bacteria are grown overnight in a series of increasing antibiotic concentrations, with or without ARM-1 present. The following day, the population from the highest tolerated concentration is re-challenged with higher concentrations of antibiotic. The median MIC is determined after each 24 hour growth (which corresponds to 7 to 10 generations) period as the lowest concentration of antibiotic at which at least 50% growth impairment is observed compared to an untreated population. MIC fold change is reported as the change in MIC from the first day of growth to each subsequent day. **c** Minimum inhibitory concentration of ARM-1 in *E. coli* and *S. aureus* after a minimum of 60 generations.

If a compound is truly effective at inhibiting a mechanism underlying resistance development, bacteria should not acquire resistance to the compound itself during exposure. To test this hypothesis, we repeatedly challenged *S. enterica* and *S. aureus* (representative gram-negative and gram-positive pathogens) with increasing concentrations of ARM-1 to measure MIC. Over the course of 60 to 70 generations (196 hours), the MIC remains constant at 400μM ARM-1 in both species (Fig. 4c, S.I. Fig. 7). Similarly, growth dynamics are not significantly impacted by sub-MIC ARM-1 treatment (S.I. Fig. 7). These observations suggest both that ARM-1 toxicity is not solely due to on-target effects and that selection pressure on any potential ARM-1 target mutations is not significantly influencing the population.

## Discussion

In addition to our discovery of ARM-1 and its impact on AMR, this work has provided significant insight into Mfd function. Our *in vitro* and *in vivo* findings show that Mfd not only requires its ATPase hydrolysis activity to remove stalled RNAPs from DNA, but that it also harbors an allosteric mechanism that is essential for this process. This is consistent with previously characterized transcription terminators, which also use allosteric mechanisms^30^. This finding makes a valuable contribution to the field and provides a new tool (ARM-1) to further study the mechanism by which Mfd removes RNAP from DNA.

The universal impact of ARM-1 on highly divergent bacteria strongly suggests that we may be able to develop a single drug that can be used against AMR development in many different pathogens. The development of ARM-1 for clinical use could provide us with one anti-evolution drug that can be used under many circumstances when AMR is a problem, regardless of the treatment or the pathogen—including those that already carry resistance mutations such as multi-drug resistant strains of *Staphylococcus aureus* or *Mycobacterium tuberculosis*.

The findings described here—the discovery of ARM-1—show that it is possible to inhibit the rise of AMR with a small molecule that targets a fundamental bacterial process. The discovery of new antibiotics directed at both new and established targets is invaluable. However, no therapies are immune to mutagenesis, the driver of adaptative evolution. When essential processes are pharmaceutically inhibited, we are essentially performing a genetic screen, selecting for mutants resistant to that selection pressure. This inevitably places all novel treatments under the threat of adaptive evolution by pathogens. The ultimate solution to protecting the efficacy of new therapeutics and limiting the development of resistant infections is to inhibit the ability of bacteria to evolve resistance during treatment. Our discovery of ARM-1 is a foundation for the development of at least one anti-evolution drug to combat the rise of AMR.

## Acknowledgments

Current and former Merrikh Lab members, especially Mark Ragheb and Maureen Thomason, the Vanderbilt Institute of Chemical Biology Molecular Design and Synthesis Center, the Vanderbilt Institute of Chemical Biology High Throughput Screening Core, VU Molecular Cell Biology Core, and Calibr at the Scripps Research Institute.

## Author Contributions and Affiliations

AEJ, HEB, AJHV, JCG, and ENS performed experiments. HM, AEJ, HEB, and JCG analyzed data. AEJ and JCG created figures. HM and AEJ wrote the manuscript. KK synthesized and validated ARM-1. HM conceived and directed the project.

## STAR Methods

### Resource Availability

#### Lead Contact

Further information and requests for resources and reagents should be directed to and will be fulfilled by the lead contact, Houra Merrikh.

#### Materials Availability

All bacterial strains and plasmids generated during this study are available upon request.

#### Data and Code Availability

All data reported in this paper will be shared by the lead contact upon request. Any additional information required to reanalyze the data reported in this paper is available from the lead contact upon request. This paper does not report any original code.

## Experimental Model Details

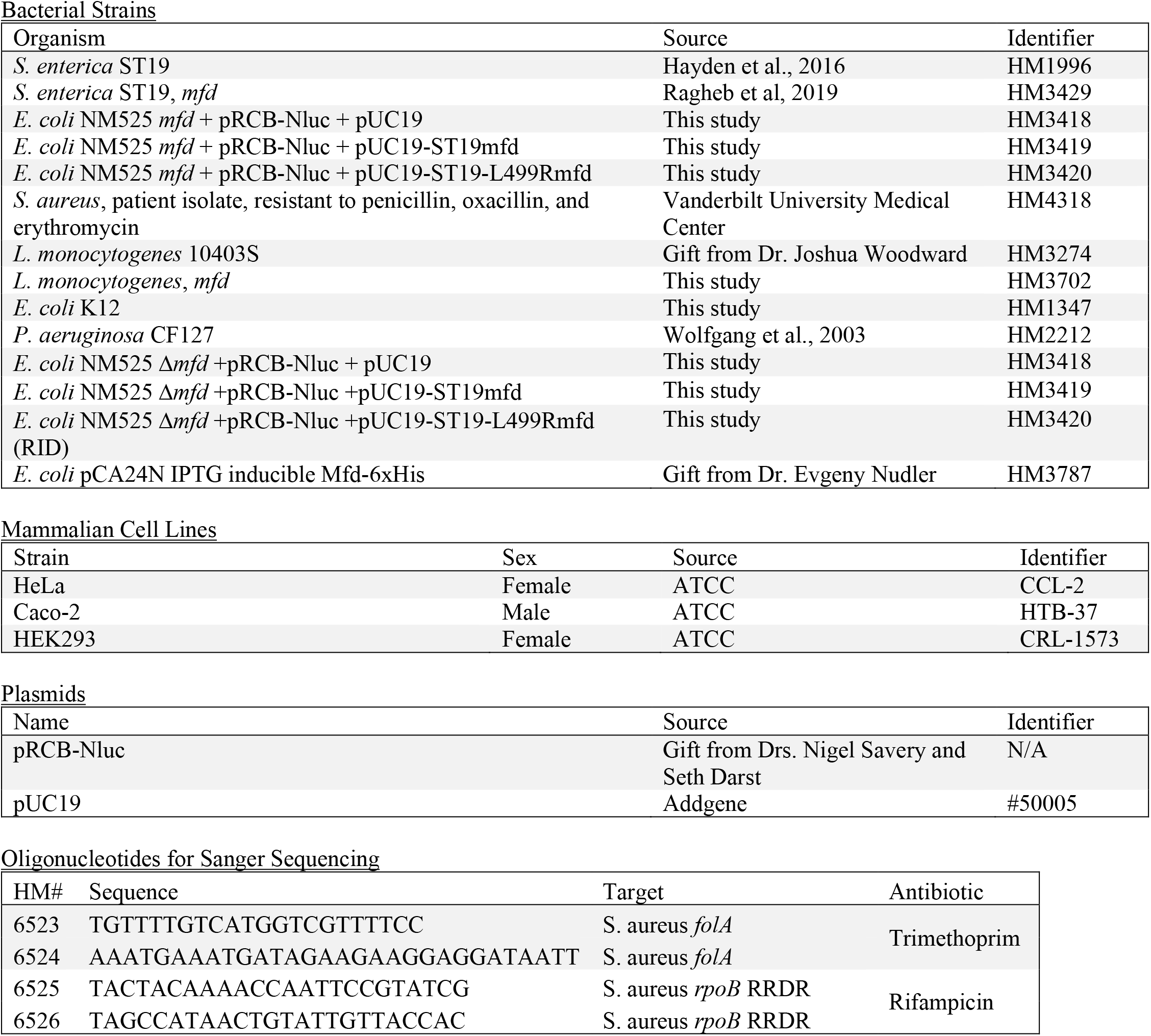
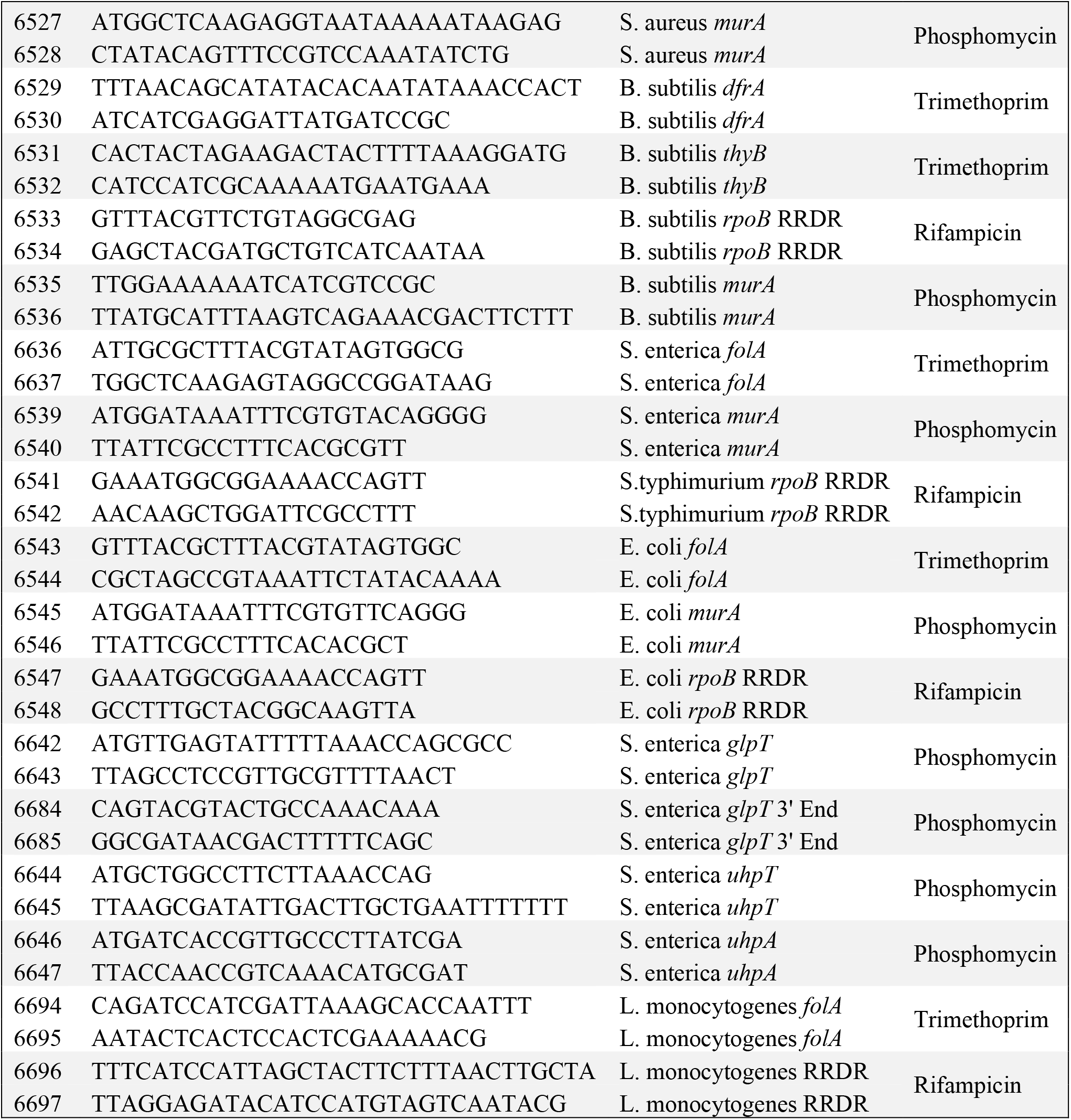

### Preparation of ARM-1 (VU001); see Supplemental Information Figs. 8-10

#### Materials

Solvents were obtained from either an MBraun MB-SPS solvent system or freshly distilled (tetrahydrofuran was distilled from sodium-benzophenone; toluene was distilled from calcium hydride and used immediately; dimethyl sulfoxide was distilled from calcium hydride and stored over 4 Å molecular sieves). Commercial reagents were used as received. The molarity of *n*-butyllithium solutions was determined by titration using diphenylacetic acid as an indicator (average of three determinations).

#### Instrumentation

Semi-preparative reverse phase HPLC was conducted on a Waters HPLC system using a Phenomenex Luna 5 μm C18(2) 100A Axia 250 × 10.00 mm column or preparative reverse phase HPLC (Gilson) using a Phenomenex Luna column (100 Å, 50 × 21.20 mm, 5 μm C18) with UV/Vis detection. Infrared spectra were obtained as thin films on NaCl plates using a Thermo Electron IR100 series instrument and are reported in terms of frequency of absorption (cm^-1^). ^1^H NMR spectra were recorded on Bruker 400, 500, or 600 MHz spectrometers and are reported relative to deuterated solvent signals. Data for ^1^H NMR spectra are reported as follows: chemical shift (δ ppm), multiplicity (s = singlet, d = doublet, t = triplet, q = quartet, p = pentet, m = multiplet, br = broad, app = apparent), coupling constants (Hz), and integration. ^13^C NMR spectra were recorded on Bruker 100, 125, or 150 MHz spectrometers and are reported relative to deuterated solvent signals. LC/MS was conducted and recorded on an Agilent Technologies 6130 Quadrupole instrument.

#### Compound preparation

*Compound 1*. (5-formylfuran-2-yl)boronic acid (1.0 g, 7.15 mmol), 4-bromo-1-iodo-2-methylbenzene (1.63 g, 5.50 mmol), sodium carbonate (2 mL, 2 M solution), and bis(triphenylphosphine)palladium(II) dichloride (193 mg, 0.28 mmol) were added to pressure vessel and dissolved in dimethoxyethane (2 mL) and ethanol (3.5 mL). The solution was sparged with argon for 5 minutes, and the reaction was then sealed and heated to 65 °C for 12 hr. The reaction was then cooled to room temperature and diluted into ethyl acetate/H_2_O (30 mL, 1:1). The organic layer was separated, and the aqueous layer was extracted with ethyl acetate (3 × 30 mL). The combined organic layers were dried over MgSO_4_, filtered, and concentrated *in vacuo*. The crude product was purified by ISCO column chromatography eluting with 0 to 40% EtOAc in hexanes to afford the brown solid title compound **1** (1.13 g, 77 % yield). ^1^H NMR (400 MHz, CDCl3) δ 9.68 (s, 1 H), 7.67 (d, *J* = 8.4 Hz 1 H), 7.46-7.41 (m, 2H), 7.33 (d, *J* = 3.6 Hz 1 H), 6.74 (d, *J* = 3.6 Hz 1 H), 2.53 (s, 3 H).

*VU001*. To a solution of compound 1 (0.5 g, 1.89 mmol) in dichloromethane (8 mL) was added N,N-dimethylpropane-1,3-diamine (0.25 g, 2.45 mmol). After stirring 2 h at room temperature, sodium triacetoxyborohydride (0.44 g, 2.08 mmol) and acetic acid (0.8 mL) were added to reaction mixture. The reaction mixture was stirred for 16 hr at room temperature, quenched with sat. sodium bicarbonate (20 mL), extracted with dichloromethane (3 × 30 mL). The combined organic phase was dried over MgSO_4_, filtered, and concentrated *in vacuo*. The crude product was purified by ISCO column chromatography eluting with 0 to 70% MeOH in dichloromethane to afford the yellow oil product (0.29 g, 44% yield). ^1^H NMR (400 MHz, CD_3_OD) δ 7.63 (d, *J* = 8.4 Hz, 1H), 7.44 (s, 1H), 7.39 (dd, d, *J* = 8.4, 1.6 Hz, 1H), 6.60 (d, *J* = 3.6 Hz, 1H), 6.42 (d, *J* = 3.6 Hz, 1H), 3.83 (s, 2H), 2.67 (t, *J* = 7.2 Hz, 2H), 2.48 (s, 3H), 2.37 (t, *J* = 7.2 Hz, 2H), 2.25 (S, 6H), 1.73 (p, *J* = 7.6 Hz, 2H); LCMS calc’d for C17H23BrN2O [M+H]^+^: 351.3 measured 352.4.

### High-Throughput Screen (Calibr)

Workflow for the primary screen is described in the main text, with additional details provided in S.I. Fig. 1. This screen was performed by Calibr (Scripps Research Institute).

### *S*. *enterica* Mfd Purification

*S. enterica* Mfd was purified by growing up HM3787 cells overnight from a single colony in LB + 20 μg/mL chloramphenicol with agitation at 37°C. The next day 1 L of fresh LB + chloramphenicol was inoculated with 10 mL of overnight culture, incubated with agitation at 37°C. at optical density of 0.3 1 mM of IPTG was added and growth was continued for another 4 hours. Cells were then centrifuged at 5,000 rpm for 10 min and washed with PBS. Cell paste was then resuspended in 30 mL HisTrap lysis buffer (50 mM NaPO_4_ pH = 8.0; 0.5 mM DTT; 0.5 M NaCl; 5 mM imidazole pH = 8.0) plus 2x Roche inhibitor proteases pills and homogenized. Cells were sonicated at midi-tip for 5 minutes at 50% amplitude with 30 second pulse intervals in an ice bath then centrifuged at 15,000 rpm for 30 minutes at 4°C. Supernatant was loaded on a His-Trap column (6 mL) in the same buffer, washed with NiWash Buffer (50 mM NaPO_4_, 300 mM NaCl, and 40 mM imidazole pH 7.4), then eluted with NiElution Buffer (50 mM NaPO_4_, 300 mM NaCl, and 150 mM imidazole pH7.4). One 15 mL fraction was collected and dialyzed overnight in a Slide-A-Lizer (Thermo Scientific) against 1000 mL of TGED Buffer (10 mM Tris-HCl pH = 8.0; 5% glycerol; 0.1 mM DTT; 0.1 mM EDTA; 50 mM NaCl). Dialysis material was centrifuged at 15,000 rpm for 30 minutes at 4°C. Using an AKTA system supernatant was loaded at 3x Heparin (1 mL) columns (GE) in TGED Buffer + 50 mM NaCl buffer at 0.6 mL/min (pressure 0.5 mPa) and eluted by the same buffer with 1 M NaCl in linear gradient from 0% to 100% in 20 column volumes. Fractions were 1 mL during loading and 1 mL during separation. Peaks with a correct molecular weight from Heparin purification were diluted to 50 mL in TGED buffer without NaCl and loaded at MonoQ 5/50 column (GE) in TGED buffer + 50 mM NaCl buffer at 2 mL/min (pressure 3.4 mPa) and eluted by the same buffer with 1 M NaCl in linear gradient from 0% to 100% in 20 column volumes. Fractions were 1 mL during loading and 0.5 mL during separation. Eluate was dialyzed overnight against 1000 mL of 10 mM RA Buffer (10 mM HEPES, 1 mM DTT, 50% glycerol, 1 mM EDTA, 500 mM KCl, and 40 mM MgCl_2_ pH 8.0) in a Slide-A-Lyser. Resulting solution was aliquoted in 0.1 mL parts and flash-frozen in liquid nitrogen before being stored at -80°C.

### Translocase Assay

HM3418 (*E. coli* NM525 Δ*mfd* +pRCB-Nluc + pUC19) and HM3419 (*E. coli* NM525 Δ*mfd* +pRCB-Nluc +pUC19-ST19mfd) were grown overnight from single colonies in the presence of antibiotic selection. Overnight cultures were back-diluted the following morning to OD600 0.05, then grown in the presence or absence of ARM-1 at 37°C until cultures reach OD600 0.50. In a 96-well plate, 80μL of cells from each sample were combined with 100μL LB and 20μL of luminescence reporter substrate (NanoLuc, Promega). Luminescence of each well quantified on a BioTek SynergyNeo plate reader.

### Binding Affinity Analysis

Microscale thermophoresis was performed on a Monolith (NanoTemper). Purified Mfd was His-tagged labeled using NanoTemper’s His-Tag Labeling Kit (MO-L018) according to manufacturer’s protocol. Serial 1:2 dilutions were made of C5 starting with an end concentration of 1 mM in PBST buffer for a total of 16 dilutions. Labeled Mfd was then added to each dilution with an end concentration of 50 nM. MST experiments were performed under default settings apart from fluorescence intensity set to 100%. The raw data was imported into Prism 9 software and nonlinear regression was used to find the K_d_.

### NADH-Coupled ATPase Assay

ATPase assays were performed as described in Kiiansita et al., 2003. All reactions were performed at 37°C in a 150μL reaction volume in a 96 well plate. Reactions were performed in repair buffer (40 mM HEPES pH 8.0, 100 mM KCl, 8 mM MgCl_2_, 4% glycerol (v/v), 5 mM DTT and 100 μg/ml BSA) supplemented with 4.4 units pyruvate kinase, 5.7 units lactate dehydrogenase, 500 μM phosphoenolpyruvate and 400 μM NADH. Mfd in a final concentration of 50nM and ARM-1 (aqueous solution, pH 8.0) in varying concentrations were added at least 15 minutes prior to starting reaction and allowed to incubate on ice. To start reaction, varying quantities of ATP were added and absorbance at 340nm was measured every 60 seconds for 1 hour in an Epoch2 microplate spectrophotometer (BioTek).

### RNAP Displacement Assay

5 ng of a ^32^P labeled, 176 bp PCR fragment containing the promoter of the ampicillin resistance gene from pDR110, a gift from David Rudner, were incubated with 1 unit of E. coli RNA Polymerase holoenzyme (NEB) for 15 mins at 37 °C. Then, NTPs were added to a final concentration of 1.7 mM (ATP) or 80 uM (UTP, GTP), as well as 80 uM ApU (Jena Bioscience), and incubated for 15 mins at 37 °C. Purified *S. enterica* ST19 Mfd (final concentration of 100 nM) that had been pre-incubated for 10 mins at 37 °C with the indicated amounts of ARM-1 (aqueous solution, pH 8.0) was added and incubated for 6 mins at 37 °C. 4 ul of the reaction were loaded into a polyacrylamide gel that had been pre-run for 45 mins at 70 V, and run for 55 mins at 150 V on ice, using 1X TBE buffer. The products were analyzed by phosphorimaging (GE Healthcare) and quantified using Image Lab 6.0.1.

#### Template Sequence

TTAGACGTCAGGTGGCACTTTTCGGGGAAATGTGCGCGGAACCCCTATTTGTTTATTTTTCTAAATACATTCAAATATGTATCCGCTCATGAGACAATAACCCTGATAAATGCTTCAATAATATTGAAAAAGGAAGAGTATGAGTATTCAACATTTCCGTGTCGCCCTTATTCCCT

### Luria-Delbruck Fluctuation Analysis

Overnight cultures of WT and *mfd* S. enterica serovar Typhimurium ST19 were grown from single colonies and back-diluted to OD_600_ 0.0005. These cultures were grown to OD_600_ = 0.8-1.0, in the presence and absence of 100μM ARM-1. 1mL of this exponential phase culture was then centrifuged at 5000rpm for 5mins at room temperature, resuspended in 100uL 1X Spizzen’s salts, and plated on LB supplemented with 80ng/mL ciprofloxacin. An additional sample of the same cultures was serially diluted an plated on LB without antibiotic selection to enumerate colony forming units (CFUs).

LB+ciprofloxacin plates were incubated at 37°C and LB plates were incubated at 30°C overnight, and all colonites counted the following morning. Mutation rates were calculated using the Ma-Sandri-Sarkar Maximum Likelihood method from a minimum of 50 biological replicates per genotype and treatment condition^36^.

### Evolution Assays

Evolution experiments were performed for the indicated strains. For *S. enterica, S. aureus, L. monocytogenes, P. aeruginosa*, and *E. coli*, overnight cultures, started from a single colony, were back diluted to OD600 = 0.005 and used to inoculate a 96-well plate. Cells were grown for 24 hours with agitation, at 37°C, in LB with a gradient of concentrations of the indicated antibiotic to select for resistance. Optical densities were subsequently measured in an Epoch2 microplate spectrophotometer (BioTek). Cultures that grew (defined by at least 50% growth relative to LB only) at the highest concentration of antibiotic were passaged into fresh LB with antibiotic in a subsequent plate. A total of 5 serial passages were performed. For *P. aeruginosa*, LB + 0.01% tween 80 was used. For all species, antibiotics were diluted 2-fold down each given row in a 96 well plate. For sequencing of resistance loci from select evolution assays, gDNA was extracted from bacterial samples using GeneJet Genomic DNA Purification Kit (ThermoFisher). Samples were sequenced by Genewiz using custom primers.

### Mutagenesis Measurements Post Epithelial Cell Infection

HELA cells were cultured in DMEM supplemented with 20% heat-inactivated FBS, 1% glutamine, and 1% penicillin/streptomycin at 37°C in 5% CO_2_. The night before infection, 1×10^7^ HeLa cells were seeded into 15cm plates and incubated overnight. A single *S. enterica* colony from strain HM1996 (WT ST19) or HM3429 (Δ*mfd*ST19) was used to inoculate an overnight culture, grown in LB with appropriate antibiotic selection at 37°C 260rpm. The next morning, the overnight bacterial culture was set back to OD_600_ 0.05 and grown to mid-exponential phase. The bacteria were then washed twice with tissue culture grade 1X PBS and resuspended in an appropriate volume of DMEM + 20% FBS + 1% glutamine. Bacteria were then applied to the HeLa cells and allowed to invade for 60 minutes at 37°C 5% CO_2_, at an MOI of 100:1. After 60 minutes of invasion, the bacteria were removed and fresh media applied to the HeLa cells. For ARM-1 treated conditions, the fresh media contained 31.25μM ARM-1. 30 minutes later, 50μg/mL gentamicin was added to the media to kill any remaining extracellular bacteria. After 8 hours of infection, HeLa cells were washed 2X with 1X PBS and lysed with 5mL 1% Triton-X-100 in water. A small portion of lysate was serially diluted and plated on LB for CFU enumeration. The remaining volume was plated on LB + 50mg/mL 5-fluorocytosine (5-FC) and grown overnight at 37°C to determine mutation frequency. Mutation frequency is reported as the ratio of number of colonies that grew on selection plates to the number of viable bacterial cells in each lysate.

### Mammalian Cell Cytotoxicity

Cytotoxicity of ARM-1 against HeLa, Caco-2, and HEK293 cells was determined using CellTox Green Cytotoxicity Assay (Promega) according to manufacturer instructions. ARM-1 was diluted in DMSO and applied to the culture such that final concentrations of DMSO were less than 0.05%. Reported toxicity is relative to a DMSO only control. Cells were incubated with ARM-1 for 24 hours prior to addition of reporter dye. Fluorescence was read on a BioTek Synergy Neo plate reader.

### Statistical Analysis

All statistical analysis was performed using GraphPad Prism 9.0.

### Figure Design

All figures were created using GraphPad Prism 9.0, Adobe Illustrator, or BioRender.

## Supplemental Information Titles and Legends

**Supplemental Information Figure 1. a** Workflow of Calibr library screening. **b** Luminescence response curve from ARM-1 performance in original screen in the presence of Mfd and active transcription. **c** Luminescence response curve from ARM-1 performance in original screen in the absence of Mfd. **d** Antibacterial activity of ARM-1 in original screen.

**Supplemental Information Figure 2**. Related to Figure 2b. NADH-Coupled ATPase assay control conditions. 100μM ARM-1 was incubated with repair assay buffer supplemented with 4.4 units pyruvate kinase, 5.7 units lactate dehydrogenase, 500 μM phosphoenolpyruvate, and 50nM Mfd, in the presence or absence of NADH and ATP.

**Supplemental Information Figure 3**. Related to Figure 2c. **a** Transcription roadblock assay, as described in Figure 2c. Each gel shows products of three independent experiments. The left four lanes of the middle gel image are the same as presented in Figure 2c, provided here for comparison. **b** Quantification of the gel images shown in **a**. Replicates A, B, C correspond to the left, middle, and right gel images, respectively. Relative density of initiation complex (IC) and elongation complex (EC) populations shown relative RNAP only condition. Analysis was performed using Image Lab 6.0.1 and summary statistics calculated using Excel. **c** Graphical representation of data shown in **b**.

**Supplemental Information Figure 4**. Related to Figure 2c. Proposed model of ARM-1 effect on interaction between Mfd and stalled transcription elongation complexes. Left panel shows proposed behavior of Mfd and RNAP in the absence of ARM-1. Increased population of initiation complexes (ICs) and decreased elongation complexes (ECs) after the addition of Mfd (Fig. 2c, *left panel, lane 3*) suggests that after Mfd displaces RNAP from the DNA template, RNAP is “recycled” and returns to the promoter to re-initiate transcription. Right panel shows proposed behavior of Mfd and RNAP in the presence of ARM-1. Under these conditions, a decrease in EC population suggests that Mfd does displace stalled RNAPs, but lack of a simultaneous increase in IC population suggests that RNAP is not effectively recycled to the promoter and does not reinitiate transcription under these conditions.

**Supplemental Information Figure 5**. Related to Figure 2d. ChIP-qPCR of rpoB enrichment at 23S rDNA. *S. enterica* ST19 of indicated genotype grown to mid exponential phase and treated with indicated concentration of ARM-1 for 2 generations prior to harvest. IP performed using 8RB13 monoclonal antibody against rpoB. Data shown representative of 3 biological replicates.

**Supplemental Information Figure 6**. Related to Figure 3. Cytotoxicity of ARM-1 against indicated mammalian cell lines. **a** Toxicity reported from original Calibr screen data against HEK293T and HepG2 cell lines. **b** Toxicity determined using Promega CellTox reagents following 8 hours of exposure of HeLa, Caco-2, and HEK293 cells to varying concentrations of ARM-1. Relative fluorescent units (RFUs) reported relative to solvent and no substrate controls.

**Supplemental Information Figure 7**. Related to Figure 4. Growth curves of indicated species in the presence of increasing concentrations of ARM-1. Precultures of each species were grown with appropriate antibiotic selection, then diluted back to OD600 0.05 in a 96 well plate. Indicated concentrations of ARM-1 were added, and plates incubated overnight at 37°C with shaking. OD measurements taken every 60s by BioTek plate reader. Data shown are representative of at least 6 biological replicates. Growth curve for *S. enterica* also confirms MIC of 400μM ARM-1 for this species, with no growth observed over the experimental timeframe.

**Supplemental Information Figure 8**. Compound 1 NMR Spectrum.

**Supplemental Information Figure 9**. VU001 H-NMR Spectrum.

**Supplemental Information Figure 10**. VU001 Liquid Chromatography with Tandem Mass Spectrometry (LCMS).

## References

1. Dadgostar P. Antimicrobial Resistance: Implications and Costs. Infect Drug Resist. 2019;12:3903–3910. doi:10.2147/IDR.S234610

2. Pribis JP, García-Villada L, Zhai Y, et al. Gamblers: An Antibiotic-Induced Evolvable Cell Subpopulation Differentiated by Reactive-Oxygen-Induced General Stress Response. Molecular Cell. 2019;74(4):785-800.e7. doi:10.1016/j.molcel.2019.02.037

3. Cirz RT, Chin JK, Andes DR, Crécy-Lagard V de, Craig WA, Romesberg FE. Inhibition of Mutation and Combating the Evolution of Antibiotic Resistance. PLOS Biology. 2005;3(6):e176. doi:10.1371/journal.pbio.0030176

4. Smith PA, Romesberg FE. Combating bacteria and drug resistance by inhibiting mechanisms of persistence and adaptation. Nat Chem Biol. 2007;3(9):549-556. doi:10.1038/nchembio.2007.27

5. Compositions and methods to reduce mutagenesis - Patent US-2006111302-A1 - PubChem. Accessed September 11, 2022. https://pubchem.ncbi.nlm.nih.gov/patent/US-2006111302-A1

6. Ragheb MN, Thomason MK, Hsu C, et al. Inhibiting the Evolution of Antibiotic Resistance. Mol Cell. 2019;73(1):157-165.e5. doi:10.1016/j.molcel.2018.10.015

7. Westblade LF, Campbell EA, Pukhrambam C, et al. Structural basis for the bacterial transcription-repair coupling factor/RNA polymerase interaction. Nucleic Acids Res. 2010;38(22):8357-8369. doi:10.1093/nar/gkq692

8. Park JS, Marr MT, Roberts JW. E. coli Transcription repair coupling factor (Mfd protein) rescues arrested complexes by promoting forward translocation. Cell. 2002;109(6):757-767. doi:10.1016/s0092-8674(02)00769-9

9. Lindsey-Boltz LA, Sancar A. The Transcription-Repair Coupling Factor Mfd Prevents and Promotes Mutagenesis in a Context-Dependent Manner. Front Mol Biosci. 2021;8:668290. doi:10.3389/fmolb.2021.668290

10. Kapoor G, Saigal S, Elongavan A. Action and resistance mechanisms of antibiotics: A guide for clinicians. J Anaesthesiol Clin Pharmacol. 2017;33(3):300-305. doi:10.4103/joacp.JOACP_349_15

11. Lambert PA. Mechanisms of antibiotic resistance in Pseudomonas aeruginosa. J R Soc Med. 2002;95(Suppl 41):22–26.

12. Tenover FC. Mechanisms of antimicrobial resistance in bacteria. Am J Med. 2006;119(6 Suppl 1):S3–10; discussion S62-70. doi:10.1016/j.amjmed.2006.03.011

13. Grundmann H, Aires-de-Sousa M, Boyce J, Tiemersma E. Emergence and resurgence of meticillin-resistant Staphylococcus aureus as a public-health threat. Lancet. 2006;368(9538):874-885. doi:10.1016/S0140-6736(06)68853-3

14. Bozdogan B, Appelbaum PC. Oxazolidinones: activity, mode of action, and mechanism of resistance. Int J Antimicrob Agents. 2004;23(2):113-119. doi:10.1016/j.ijantimicag.2003.11.003

15. Hiramatsu K, Cui L, Kuroda M, Ito T. The emergence and evolution of methicillin-resistant Staphylococcus aureus. Trends Microbiol. 2001;9(10):486-493. doi:10.1016/s0966-842x(01)02175-8

16. Alekshun MN, Levy SB. Molecular mechanisms of antibacterial multidrug resistance. Cell. 2007;128(6):1037-1050. doi:10.1016/j.cell.2007.03.004

17. Kim YH, Cha CJ, Cerniglia CE. Purification and characterization of an erythromycin esterase from an erythromycin-resistant Pseudomonas sp. FEMS Microbiol Lett. 2002;210(2):239-244. doi:10.1111/j.1574-6968.2002.tb11187.x

18. Chambers AL, Smith AJ, Savery NJ. A DNA translocation motif in the bacterial transcription–repair coupling factor, Mfd. Nucleic Acids Res. 2003;31(22):6409-6418. doi:10.1093/nar/gkg868

19. Du M, Kodner S, Bai L. Enhancement of LacI binding in vivo. Nucleic Acids Res. 2019;47(18):9609-9618. doi:10.1093/nar/gkz698

20. Brugger C, Zhang C, Suhanovsky MM, et al. Molecular determinants for dsDNA translocation by the transcription-repair coupling and evolvability factor Mfd. Nat Commun. 2020;11(1):3740. doi:10.1038/s41467-020-17457-1

21. Selby CP, Sancar A. Molecular mechanism of transcription-repair coupling. Science. 1993;260(5104):53-58. doi:10.1126/science.8465200

22. Kang JY, Llewellyn E, Chen J, et al. Structural basis for transcription complex disruption by the Mfd translocase. Elife. 2021;10:e62117. doi:10.7554/eLife.62117

23. Smith AJ, Pernstich C, Savery NJ. Multipartite control of the DNA translocase, Mfd. Nucleic Acids Res. 2012;40(20):10408-10416. doi:10.1093/nar/gks775

24. Carter RH, Demidenko AA, Hattingh-Willis S, Rothman-Denes LB. Phage N4 RNA polymerase II recruitment to DNA by a single-stranded DNA-binding protein. Genes Dev. 2003;17(18):2334-2345. doi:10.1101/gad.1121403

25. Smith AJ, Savery NJ. RNA polymerase mutants defective in the initiation of transcription-coupled DNA repair. Nucleic Acids Research. 2005;33(2):755-764. doi:10.1093/nar/gki225

26. Deaconescu AM, Chambers AL, Smith AJ, et al. Structural Basis for Bacterial Transcription-Coupled DNA Repair. Cell. 2006;124(3):507-520. doi:10.1016/j.cell.2005.11.045

27. Smith AJ, Szczelkun MD, Savery NJ. Controlling the motor activity of a transcription-repair coupling factor: autoinhibition and the role of RNA polymerase. Nucleic Acids Res. 2007;35(6):1802-1811. doi:10.1093/nar/gkm019

28. Song E, Uhm H, Munasingha PR, et al. Rho-dependent transcription termination proceeds via three routes. Nat Commun. 2022;13(1):1663. doi:10.1038/s41467-022-29321-5

29. Hao Z, Svetlov V, Nudler E. Rho-dependent transcription termination: a revisionist view. Transcription. 2021;12(4):171-181. doi:10.1080/21541264.2021.1991773

30. Roberts JW. Mechanisms of Bacterial Transcription Termination. J Mol Biol. 2019;431(20):4030-4039. doi:10.1016/j.jmb.2019.04.003

31. Ragheb MN, Merrikh C, Browning K, Merrikh H. Mfd regulates RNA polymerase association with hard-to-transcribe regions in vivo, especially those with structured RNAs. Proc Natl Acad Sci U S A. 2021;118(1). doi:10.1073/pnas.2008498118

32. Richardson AR, Soliven KC, Castor ME, Barnes PD, Libby SJ, Fang FC. The Base Excision Repair system of Salmonella enterica serovar typhimurium counteracts DNA damage by host nitric oxide. PLoS Pathog. 2009;5(5):e1000451. doi:10.1371/journal.ppat.1000451

33. Barlow M, Hall BG. Experimental prediction of the natural evolution of antibiotic resistance. Genetics. 2003;163(4):1237-1241.

34. Hayden HS, Matamouros S, Hager KR, et al. Genomic Analysis of Salmonella enterica Serovar Typhimurium Characterizes Strain Diversity for Recent U.S. Salmonellosis Cases and Identifies Mutations Linked to Loss of Fitness under Nitrosative and Oxidative Stress. mBio. 2016;7(2):e00154–16. doi:10.1128/mBio.00154-16

35. Kiianitsa K, Solinger JA, Heyer WD. NADH-coupled microplate photometric assay for kinetic studies of ATP-hydrolyzing enzymes with low and high specific activities. Anal Biochem. 2003;321(2):266-271. doi:10.1016/s0003-2697(03)00461-5

36. Hall BM, Ma CX, Liang P, Singh KK. Fluctuation analysis CalculatOR: a web tool for the determination of mutation rate using Luria-Delbruck fluctuation analysis. Bioinformatics. 2009;25(12):1564-1565. doi:10.1093/bioinformatics/btp253

