## Supplementary data for "A small molecule that inhibits the evolution of antibiotic resistance"

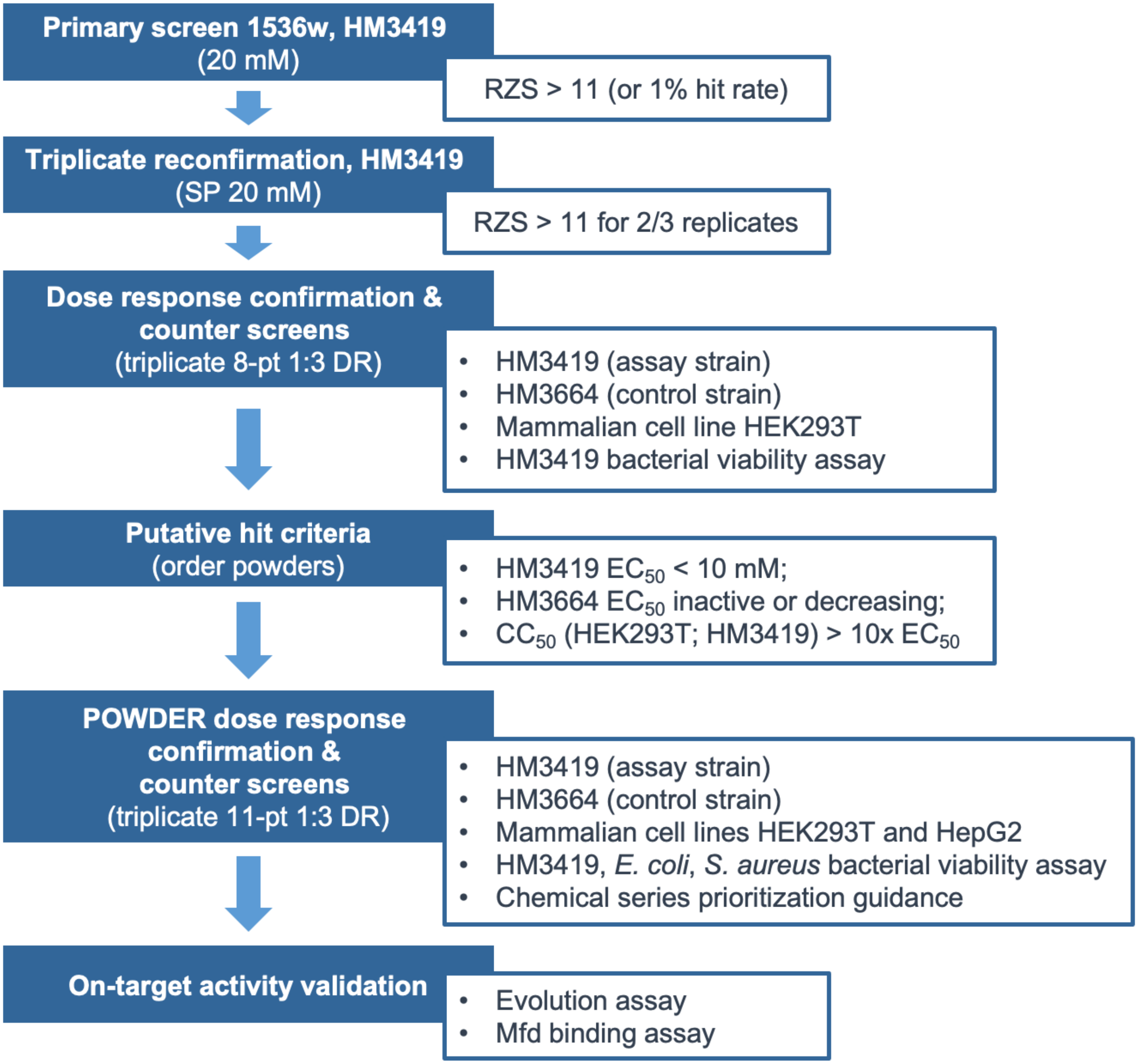


**Supplemental Information Figure 1. a** Workflow of Calibr library screening. **b** Luminescence response curve from ARM-1 performance in original screen in the presence of Mfd and active transcription. **c** Luminescence response curve from ARM-1 performance in original screen in the absence of Mfd. **d** Antibacterial activity of ARM-1 in original screen.

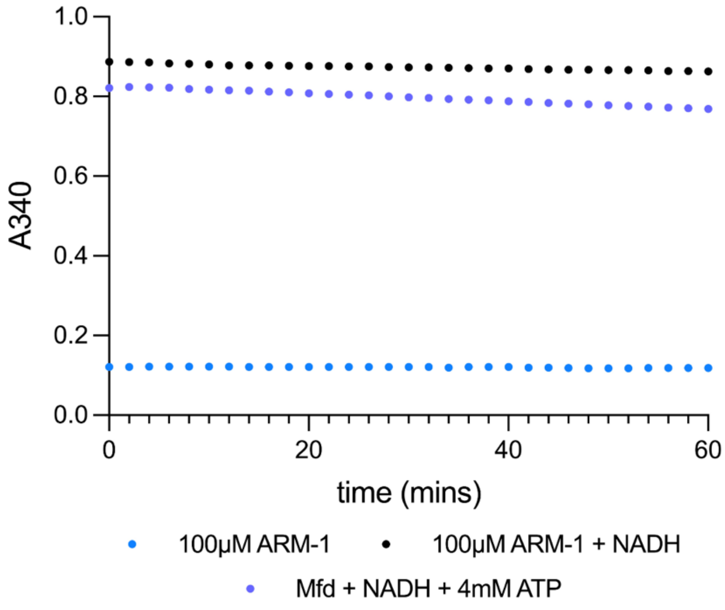


**Supplemental Information Figure 2.** Related to Figure 2b. NADH-Coupled ATPase assay control conditions. 100µM ARM-1 was incubated with repair assay buffer supplemented with 4.4 units pyruvate kinase, 5.7 units lactate dehydrogenase, 500 µM phosphoenolpyruvate, and 50nM Mfd, in the presence or absence of NADH and ATP.

**a**

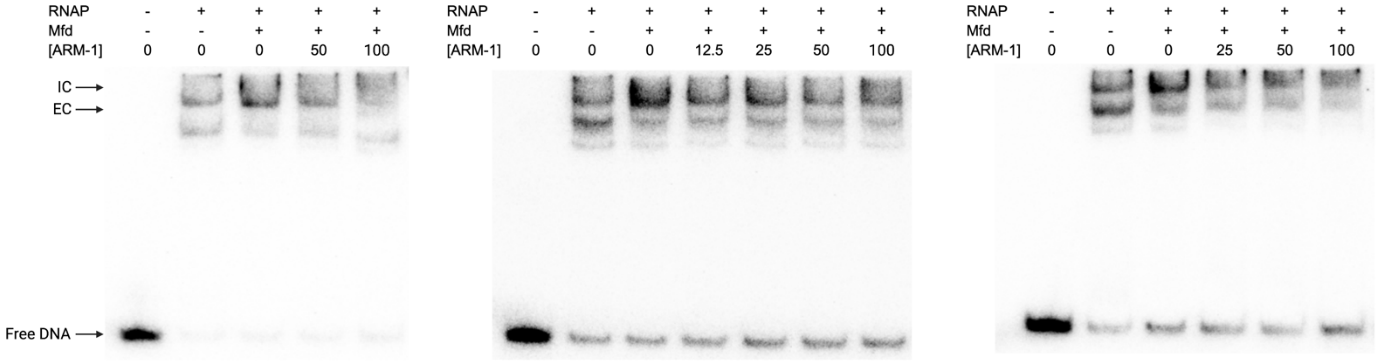


**b**

| RNAP | Mfd | ARM-1 (μM) | Band | A | B | C | Mean | Std. Dev. | EC/IC |
|---|---|---|---|---|---|---|---|---|---|
| + | - | 0 | IC | 1.00 | 1.00 | 1.00 | 1.00 | | |
| | | | EC | 1.00 | 1.00 | 1.00 | 1.00 | | 1.00 |
| + | + | 0 | IC | 1.41 | 1.76 | 2.00 | 1.72 | 0.30 | |
| | | | EC | 0.24 | 0.41 | 0.55 | 0.40 | 0.16 | 0.23 |
| + | + | 12.5 | IC | | 1.54 | 1.47 | 1.51 | 0.05 | |
| | | | EC | | 0.45 | 0.81 | 0.63 | 0.25 | 0.42 |
| + | + | 25 | IC | | | 1.45 | 1.45 | | |
| | | | EC | | | 0.80 | 0.80 | | 0.55 |
| + | + | 50 | IC | 1.29 | 1.57 | 1.27 | 1.38 | 0.17 | |
| | | | EC | 0.45 | 0.53 | 0.86 | 0.61 | 0.22 | 0.45 |
| + | + | 100 | IC | 1.13 | 1.21 | 1.37 | 1.24 | 0.12 | |
| | | | EC | 0.76 | 0.32 | 0.77 | 0.62 | 0.25 | 0.50 |

**c**

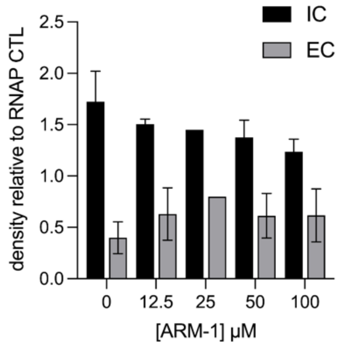


**Supplemental Information Figure 3.** Related to Figure 2c. **a** Transcription roadblock assay, as described in Figure 2c. Each gel shows products of three independent experiments. The left four lanes of the middle gel image are the same as presented in Figure 2c, provided here for comparison. **b** Quantification of the gel images shown in **a**. Replicates A, B, C correspond to the left, middle, and right gel images, respectively. Relative density of initiation complex (IC) and elongation complex (EC) populations shown relative RNAP only condition. Analysis was performed using Image Lab 6.0.1 and summary statistics calculated using Excel. **c** Graphical representation of data shown in **b**.

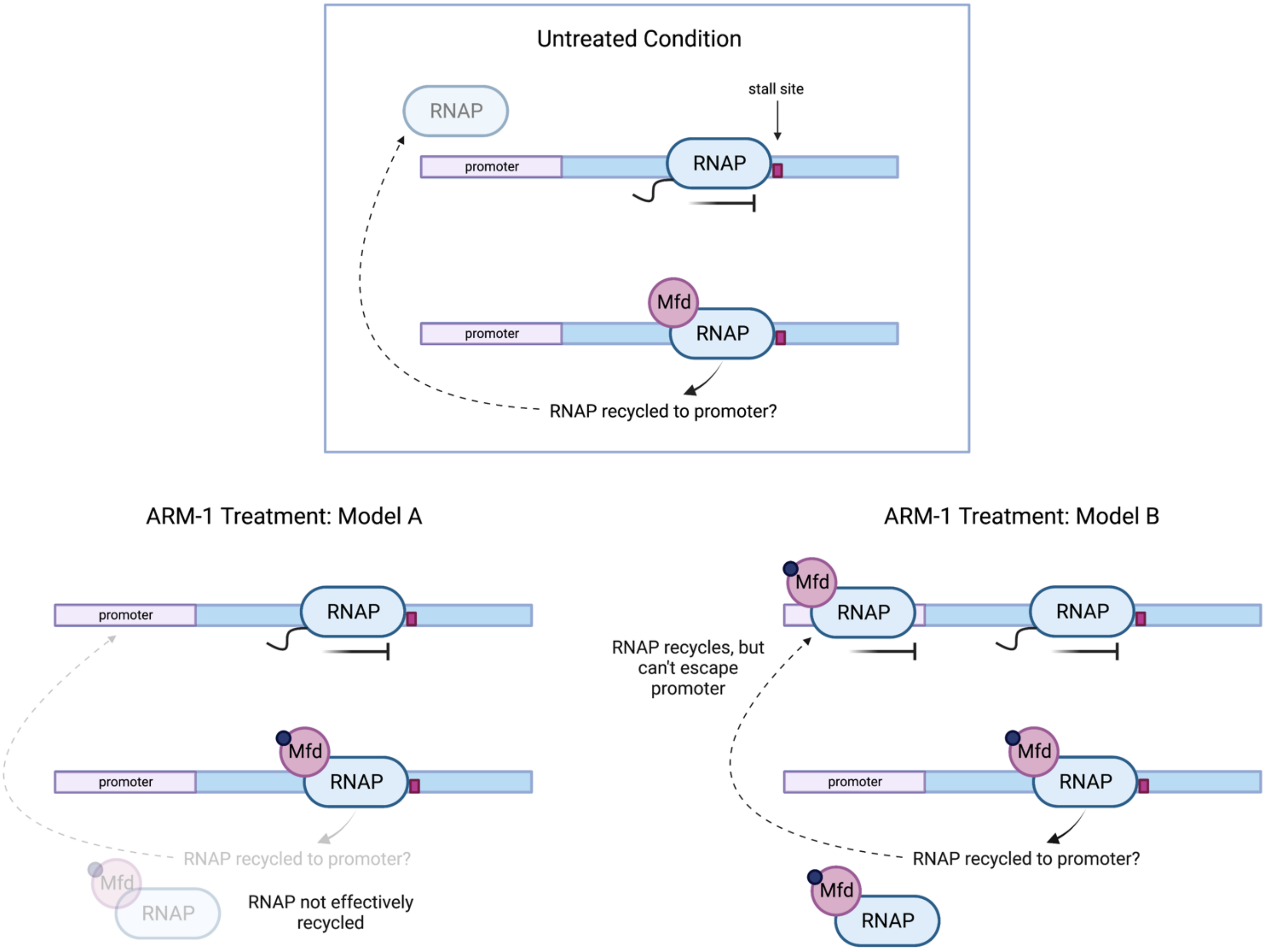


**Supplemental Information Figure 4.** Related to Figure 2c. Proposed model of ARM-1 effect on interaction between Mfd and stalled transcription elongation complexes. Left panel shows proposed behavior of Mfd and RNAP in the absence of ARM-1. Increased population of initiation complexes (ICs) and decreased elongation complexes (ECs) after the addition of Mfd (Fig. 2c, *left panel, lane 3*) suggests that after Mfd displaces RNAP from the DNA template, RNAP is "recycled" and returns to the promoter to re-initiate transcription. Right panel shows proposed behavior of Mfd and RNAP in the presence of ARM-1. Under these conditions, a decrease in EC population suggests that Mfd does displace stalled RNAPs, but lack of a simultaneous increase in IC population suggests that RNAP is not effectively recycled to the promoter and does not reinitiate transcription under these conditions.

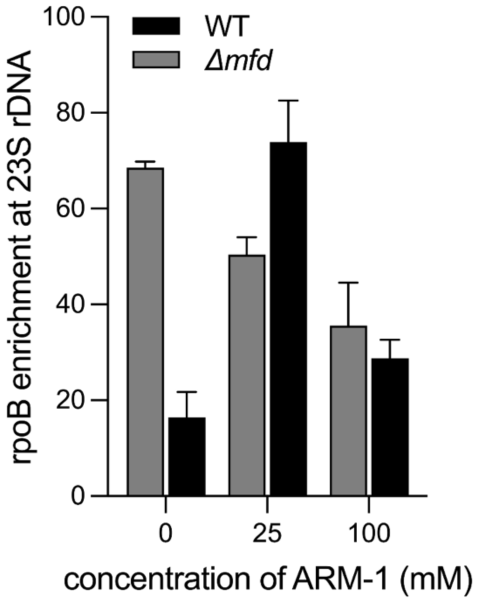


**Supplemental Information Figure 5.** Related to Figure 2d. ChIP-qPCR of rpoB enrichment at 23S rDNA. *S. enterica* ST19 of indicated genotype grown to mid exponential phase and treated with indicated concentration of ARM-1 for 2 generations prior to harvest. IP performed using 8RB13 monoclonal antibody against rpoB. Data shown representative of 3 biological replicates.

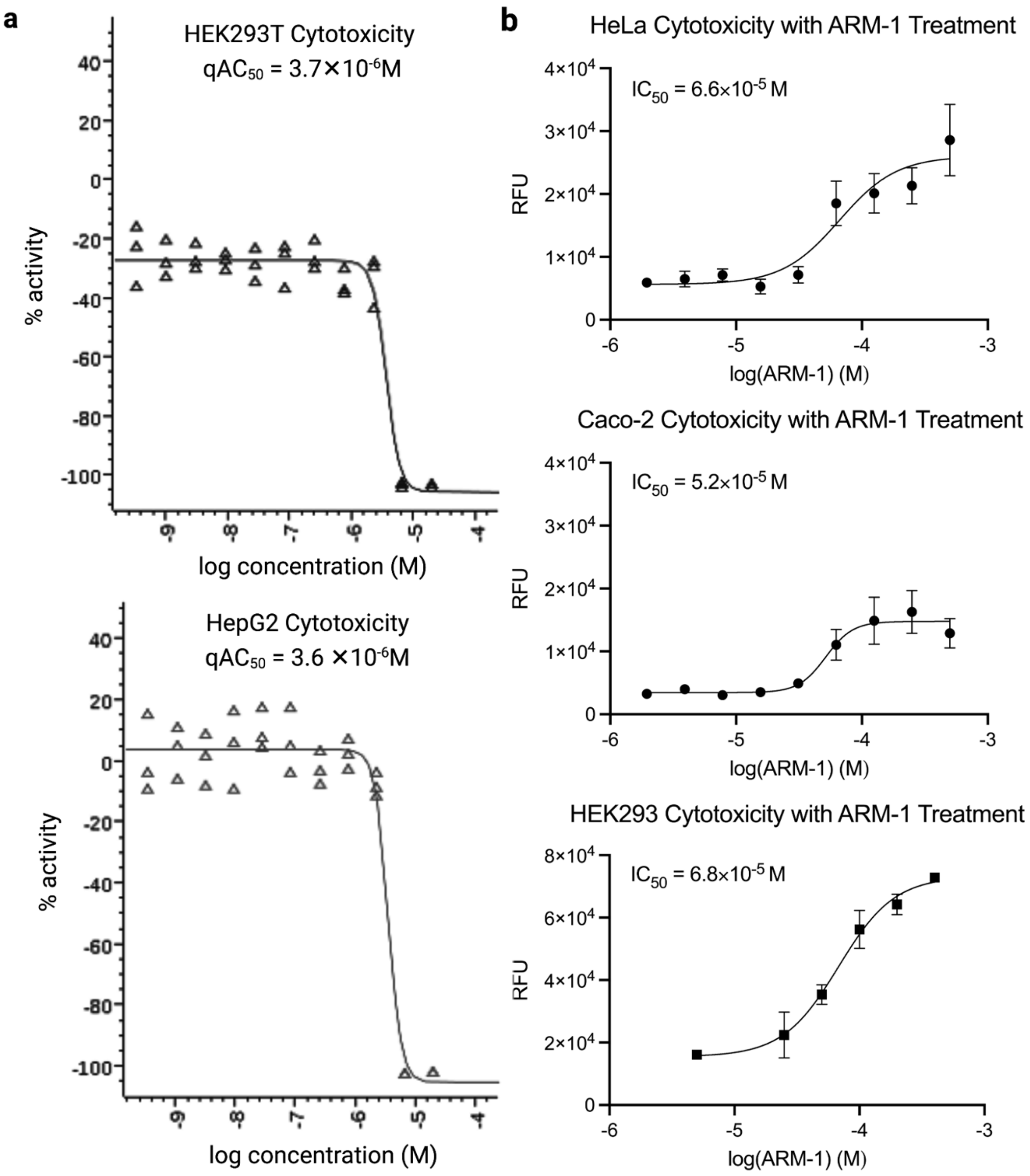


**Supplemental Information Figure 6.** Related to Figure 3. Cytotoxicity of ARM-1 against indicated mammalian cell lines. **a** Toxicity reported from original Calibr screen data against HEK293T and HepG2 cell lines. **b** Toxicity determined using Promega CellTox reagents following 8 hours of exposure of HeLa, Caco-2, and HEK293 cells to varying concentrations of ARM-1. Relative fluorescent units (RFUs) reported relative to solvent and no substrate controls.

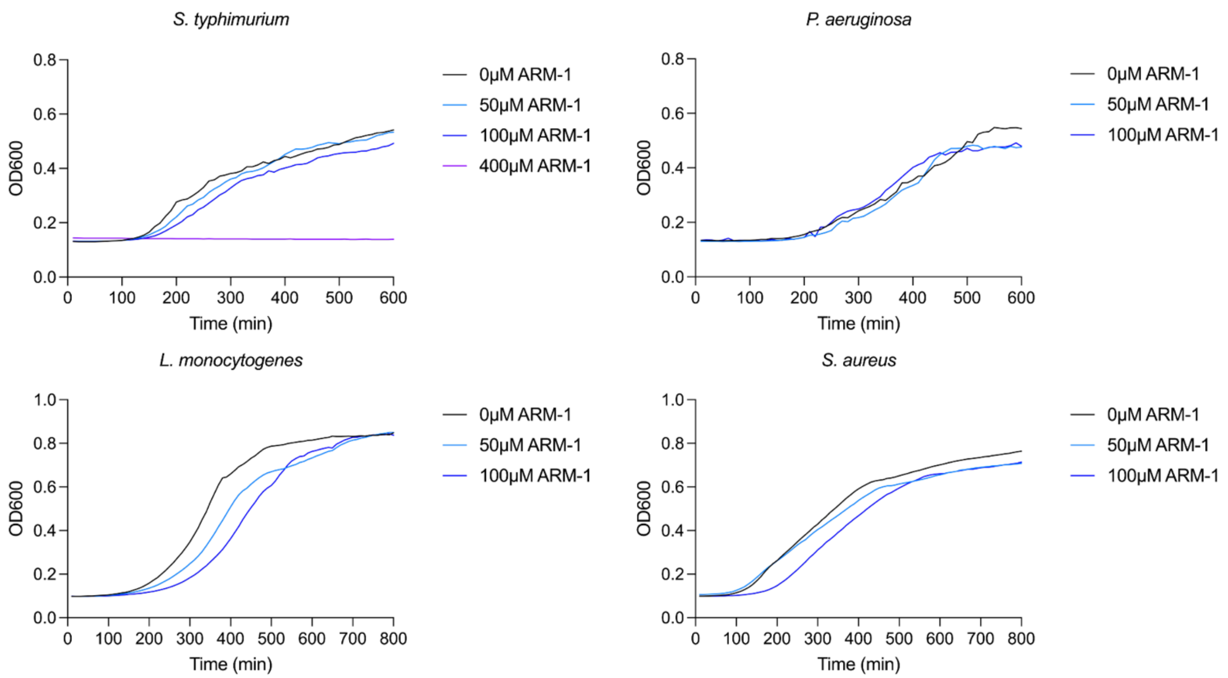


**Supplemental Information Figure 7.** Related to Figure 4. Growth curves of indicated species in the presence of increasing concentrations of ARM-1. Precultures of each species were grown with appropriate antibiotic selection, then diluted back to OD600 0.05 in a 96 well plate. Indicated concentrations of ARM-1 were added, and plates incubated overnight at 37°C with shaking. OD measurements taken every 60s by BioTek plate reader. Data shown are representative of at least 6 biological replicates. Growth curve for *S. enterica* also confirms MIC of 400µM ARM-1 for this species, with no growth observed over the experimental timeframe.

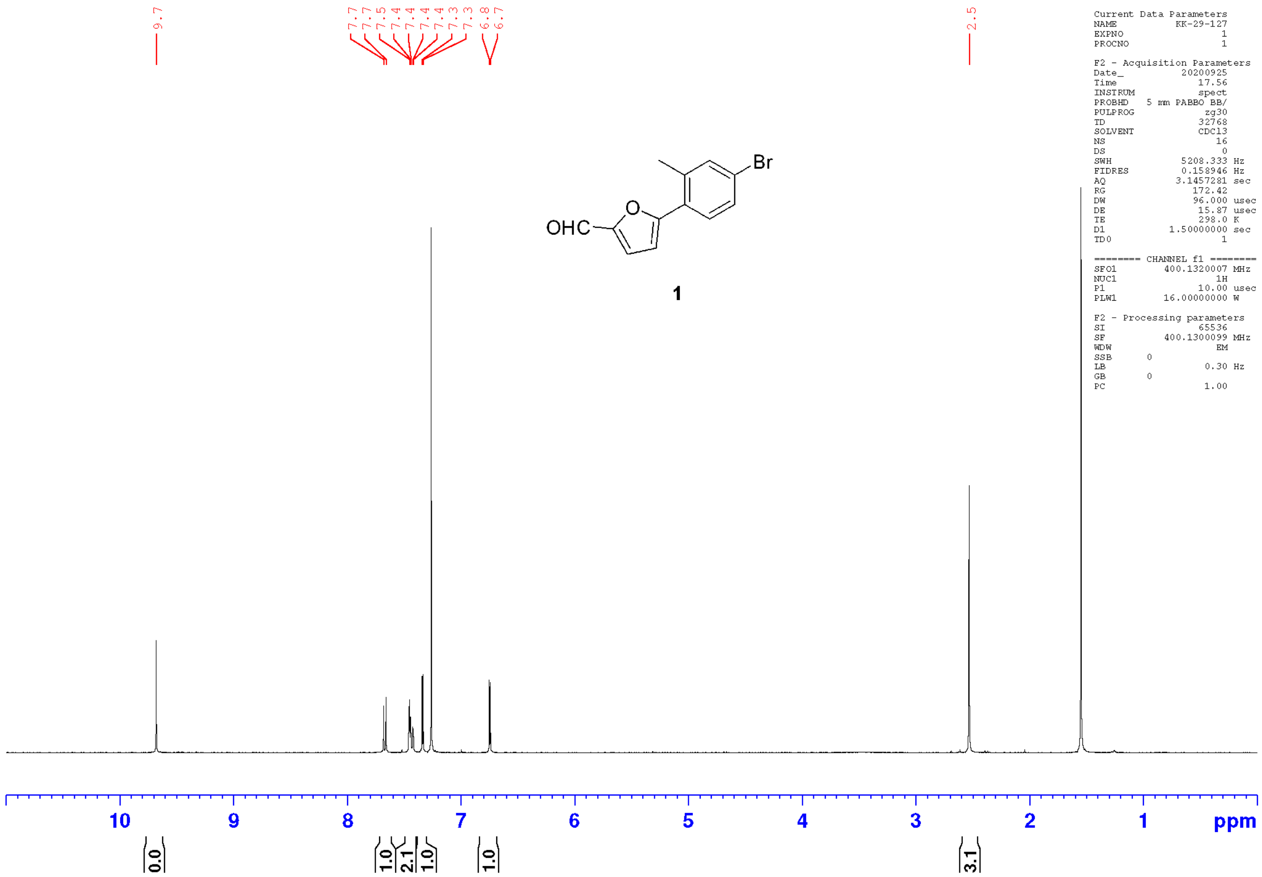

**Supplemental Information Figure 8.** Compound 1 NMR Spectrum.

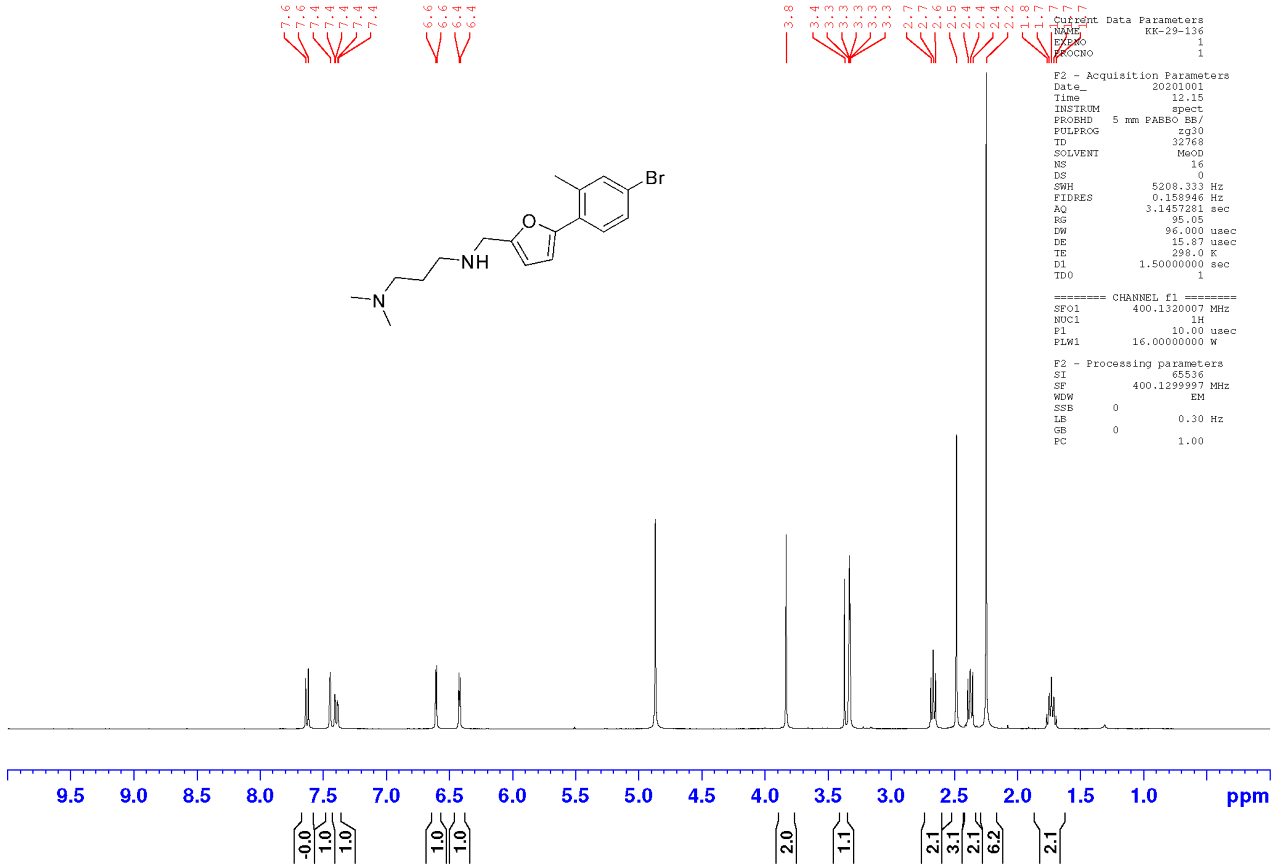


**Supplemental Information Figure 9.** VU001 H-NMR Spectrum.

**VCNDD/Syncore MRB4-12410 Agilent 1200 LCMS**

**Sample Name:** KK-29-136-P2
**User:** kimk3

**Acquired:** 9/30/2020 12:15 PM
**Filename:** C:\LCMS\DATA\KWANGHO_KIM\09-20\KK-29-136-P211-12410 SYNCORE LCMS-.D

**Method** 1 MIN.M
**Location:** 1,1:A,9

**Description:** Easy-Access Method: '1 Minute' ,351.00

| | | Area % | Area % | Area % | | Area Abs | Area Abs | Area Abs |
|---|---|---|---|---|---|---|---|---|
| Peak # | Time | UV215 | UV254 | TIC(+) | BPM | UV215 | UV254 | TIC(+) |
| 1 | 0.112 | 0.0 | 19.0 | 2.3 | 214.2 | 0 | 103.26 | 96822.67 |
| 2 | 0.841 | 100.0 | 81.0 | 97.7 | 351.3 | 641.83 | 441.29 | 4030277.75 |
| 3 | 0.919 | 0.0 | 0.0 | 0.0 | 351.3 | 0 | 0 | 0 |

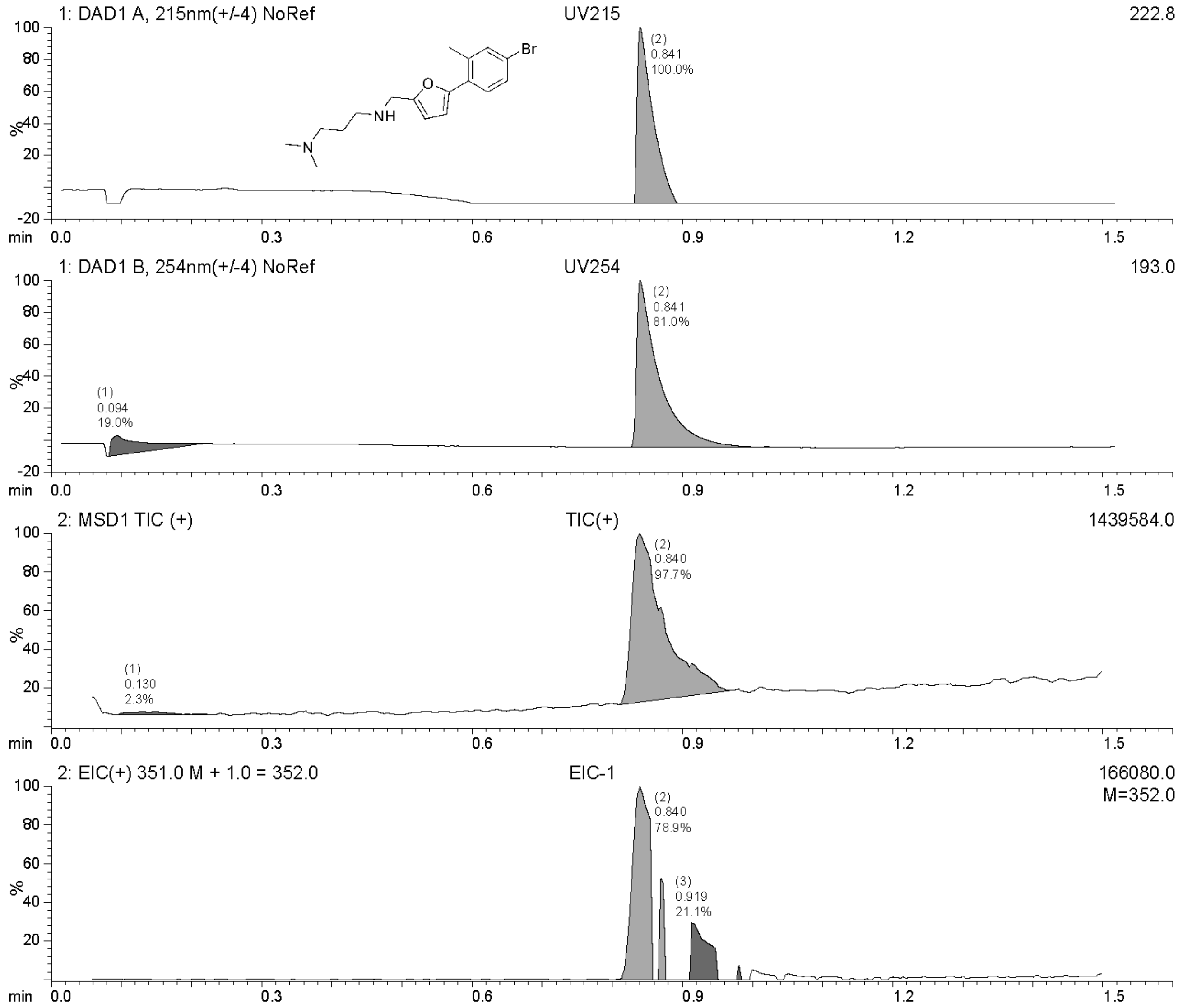

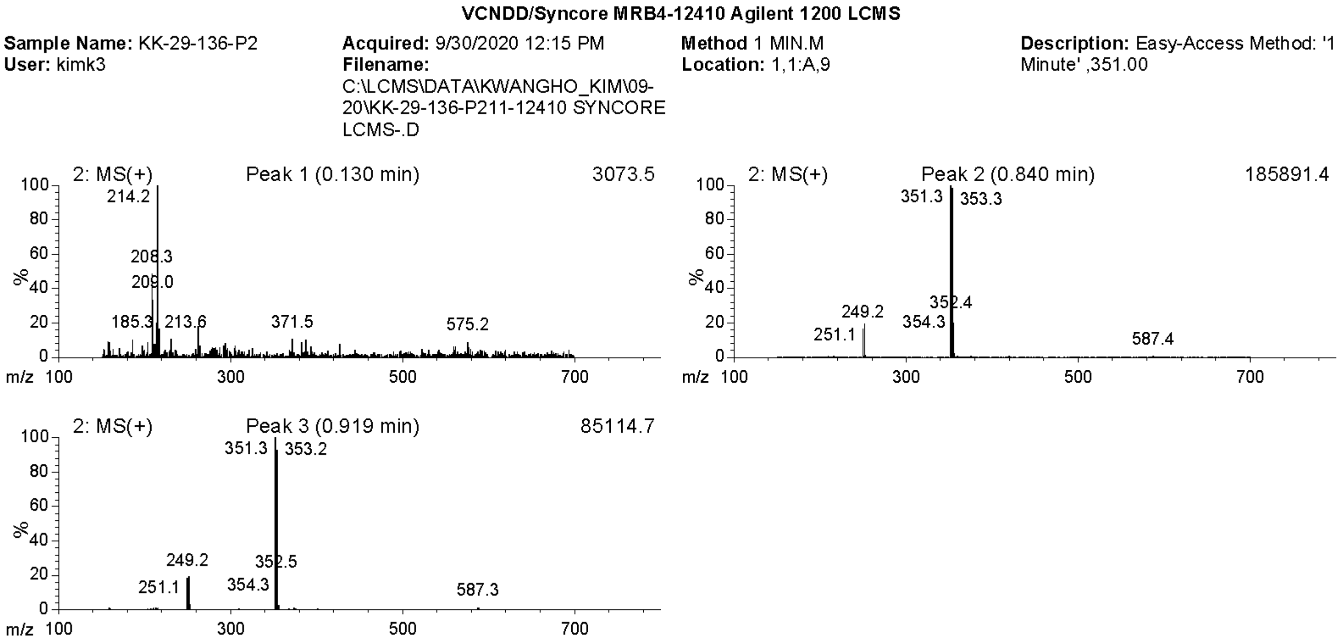


**Supplemental Information Figure 10.** VU001 Liquid Chromatography with Tandem Mass Spectrometry (LCMS).
